# Sustained ErbB activation causes demyelination and hypomyelination by driving necroptosis of mature oligodendrocytes and apoptosis of oligodendrocyte precursor cells

**DOI:** 10.1101/2020.11.10.377226

**Authors:** Xu Hu, Guanxiu Xiao, Li He, Xiaojie Niu, Huashun Li, Tianjie Lou, Qianqian Hu, Youguang Yang, Qi Xu, Zhengdong Wei, Mengsheng Qiu, Kenji F. Tanaka, Ying Shen, Yanmei Tao

## Abstract

Oligodendrocytes are vulnerable to genetic and environmental insults and its injury leads to demyelinating diseases. The roles of ErbB receptors in maintaining the CNS myelin integrity are largely unknown. Here we overactivate ErbB receptors that mediate signaling of either neuregulin or EGF family growth factors and found their synergistic activation caused deleterious outcomes in white matter. Sustained ErbB activation induced by the tetracycline-dependent mouse tool *Plp*-tTA resulted in demyelination, axonal degeneration, oligodendrocyte precursor cell (OPC) proliferation, astrogliosis, and microgliosis in white matter. Moreover, there was hypermyelination prior to these inflammatory pathological events. In contrast, sustained ErbB activation induced by another tetracycline-dependent mouse tool *Sox10*^+/rtTA^ caused hypomyelination in the corpus callosum and optic nerve, which appeared to be a developmental deficit and did not associate with OPC regeneration, astrogliosis, or microgliosis. By tracing the differentiation states of cells expressing tTA/rtTA-dependent transgene or pulse-labeled reporter proteins *in vitro* and *in vivo*, we found that *Plp*-tTA targeted mainly mature oligodendrocytes (MOs), whereas *Sox10*^+/rtTA^ targeted OPCs and newly-formed oligodendrocytes. The distinct phenotypes of mice with ErbB overactivation induced by *Plp*-tTA and *Sox10*^+/rtTA^ consolidated their non-overlapping targeting preferences in the oligodendrocyte lineage, and enabled us to demonstrate that ErbB overactivation in MOs induced necroptosis that caused inflammatory demyelination, whereas in OPCs induced apoptosis that caused non-inflammatory hypomyelination. Early interference with aberrant ErbB activation ceased oligodendrocyte deaths and restored myelin development in both mice. This study suggests that aberrant ErbB activation is an upstream pathogenetic mechanism of demyelinating diseases, providing a potential therapeutic target.

**Significance statement:** Primary oligodendropathy is one of the etiological mechanisms for multiple sclerosis, and oligodendrocyte necroptosis is a pathological hallmark in the disease. Moreover, the demyelinating disease is now a broad concept that embraces schizophrenia, in which white matter lesions are an emerging feature. ErbB overactivation has been implicated in schizophrenia by genetic analysis and postmortem studies. This study suggests the etiological implications of ErbB overactivation in myelin pathogenesis and elucidates the pathogenetic mechanisms.

## Introduction

Oligodendrocytes form myelin to speed up the axonal impulse conduction while supporting neuronal function and integrity in the central nervous system (CNS). White matter is expanded, with a growth peak during juvenile to adolescence, by oligodendrocyte precursor cell (OPC) proliferation. Terminally differentiated OPCs differentiate into newly-formed oligodendrocytes (NFOs) under stringent regulation by extrinsic and intrinsic signals (Bergles and Richardson, 2015; Emery and Lu, 2015). NFOs progress from pre-myelinating oligodendrocytes to newly myelinating oligodendrocytes. Myelinating oligodendrocytes effectively generate myelin sheaths in a short time window. After myelin formation, mature oligodendrocytes (MOs) are extremely stable in maintaining myelin sheaths (Watkins et al., 2008; Czopka et al., 2013; Xiao et al., 2016; Tripathi et al., 2017; Hughes et al., 2018). However, a plethora of factors, including genetic alteration, inflammatory mediators, ischemia, trauma, viruses, and toxins, can damage oligodendrocytes and result in demyelinating diseases (Bradl and Lassmann, 2010). Understanding the pathogenetic mechanisms is essential for developing strategies to combat myelin loss and promote myelin restoration.

Intracellular signaling pathways, such as MAPK, Akt, and mTOR, are activated to promote oligodendrocyte differentiation, survival, and myelination (Flores et al., 2000; Flores et al., 2008; Ishii et al., 2012; Bercury et al., 2014). Tyrosine kinase receptors ErbB(1-4), mediating abovementioned signal transduction pathways, are known to be fundamental to various neural developmental events (Mei and Nave, 2014). Numerous growth factors are able to activate ErbB receptors, including the neuregulin (NRG) family (NRG1-NRG6) and the epidermal growth factor (EGF) family (EGF, HB-EGF, TGFα, amphiregulin, epiregulin, β-cellulin, and epigen). Among them, most of the EGF family ligands bind to EGFR only, and the NRG family ligands bind to ErbB3 and ErbB4, except that HB-EGF, β-cellulin, and epiregulin can bind both EGFR and ErbB4 (Mei and Nave, 2014). Ligand binding stimulates ErbB receptors to dimerize with one another or preferentially with ErbB2, activating multiple downstream signaling pathways (Mei and Nave, 2014).

Although it is well-established that ErbB2 and ErbB3 are crucial to peripheral myelin development (Nave and Salzer, 2006), the roles of ErbB receptors in oligodendrocytes have long been debated. For example, ErbB3 has been reported to be dispensable for oligodendrocyte development (Schmucker et al., 2003), and ErbB3/ErbB4 double knockout from embryonic age does not result in CNS myelin alteration (Brinkmann et al., 2008). However, there is another report that ErbB3 depletion in oligodendrocytes from postnatal day 19 (P19) results in adult hypomyelination (Makinodan et al., 2012). Studies on oligodendrocyte-specific knock-out mice have validated the expression of ErbB3 and ErbB4 in oligodendrocytes (Brinkmann et al., 2008), while phosphorylated EGFR is detected in oligodendrocytes by immunostaining (Palazuelos et al., 2014). Despites the controversial results from loss-of-function studies, overexpressing NRG1 Type I or Type III in neurons, or hEGFR in oligodendrocytes (*CNP*-hEGFR), promotes the CNS myelination (Aguirre et al., 2007; Brinkmann et al., 2008). Interestingly, transgenic mice *CNP*-hEGFR increases the numbers of myelinated axons but not myelin thickness (Aguirre et al., 2007), whereas NRG1-overexpressing mice exhibit hypermyelination of individual axons (Brinkmann et al., 2008). These results suggest that EGF-ErbB and NRG-ErbB receptors play different roles in oligodendrocytes. However, the possible synergistic effects of overactivating NRG-ErbB and EGF-ErbB receptors in oligodendrocytes have not been investigated.

To understand the consequences of co-activating ErbB receptors in oligodendrocytes, we employed a pan-ErbB strategy and activated endogenous ErbB receptors mediating either NRG or EGF signaling in oligodendrocytes. The results demonstrated that overactivating ErbB receptors in oligodendrocytes caused deleterious outcomes in white matter. Further, with the finding that the two inducible mouse tools, *Plp*-tTA and *Sox10*^+/rtTA^, differentially targeted MO and OPC-NFO stages, we demonstrated that ErbB overactivation induced primary MO necroptosis and OPC apoptosis, causing inflammatory demyelination and non-inflammatory hypomyelination, respectively. Early interference with the aberrant activation of ErbB signaling was effective to cease these pathological progressions and restore oligodendrocyte numbers and myelin development.

## Materials and Methods

### Animals

*Plp*-tTA transgenic mice were from the RIKEN BioResource Center (Stock No. RBRC05446). Transgenic mice *TRE*-ErbB2^V664E^ (Stock No. 010577) was from the Jackson Laboratory. Sox10*^+/^*^rtTA^ mice were from Dr. Michael Wegner (University Erlangen-Nurnberg, Germany). Unless otherwise indicated, mice were housed in a room with a 12-hour light/dark cycle with access to food and water *ad libitum*. For biochemical and histological experiments, *Plp*-tTA;*TRE*-ErbB2^V664E^ (*Plp*-ErbB2^V664E^) and *Sox10^+/^*^rtTA^;*TRE*-ErbB2^V664E^ (*Sox10*-ErbB2^V664E^) mice with either sex and their littermate control mice with matched sex were used. Animal experiments were approved by the Institutional Animal Care and Use Committee of the Hangzhou Normal University. For genotyping, the following primers were used: for *Plp*-tTA (630bp), PLPU-604 5’-TTT CCC ATG GTC TCC CTT GAG CTT, mtTA24L 5’-CGG AGT TGA TCA CCT TGG ACT TGT; for Sox10*^+/^*^rtTA^ (618bp), sox10-rtTA1 5’-CTA GGC TGT CAG AGC AGA CGA, sox10-rtTA2 5’-CTC CAC CTC TGA TAG GT CTT G; for *TRE*-ErbB2^V664E^ (625bp), 9707 5’-AGC AGA GCT CGT TTA GTG, 9708 5’-GGA GGC GGC GAC ATT GTC.

### Tet-Off or Tet-On treatment of mice

Mice with Tet-system contain genes of tetracycline-controlled transcriptional activator (tTA) or reverse tetracycline-controlled transcriptional activator (rtTA) driven by cell-specific promoters. When fed with Dox, these mice are able to switch on or off expression of a gene under the control of *tetracycline-responsive element* (*TRE*), specifically in rtTA- or tTA-expressing cells, which are called ‘Tet-on’ or ‘Tet-off’, respectively. The offspring of *Sox10*^+/rtTA^ during the indicated periods were fed with Dox (2 mg/mL × 10 mL/day from P21 to indicated test day) in drinking water to induce the expression of ErbB2^V664E^ in *Sox10*-ErbB2^V664E^ mice (Tet-On). For the offspring of *Plp*-tTA, Dox was given (Tet-off) from the embryonic day (through pregnant mothers) to their weaning day at P21 to inhibit the expression of ErbB2^V664E^ during this period in *Plp*-ErbB2^V664E^ mice (0.5 mg/mL × 10 mL/day of Dox before P21). Water bottles were wrapped with foil to protect Dox from light. All used littermate control mice were treated the same.

### Stereotaxic injection of AAV viruses

pAAV-*TRE*-EYFP plasmids (Addgene) were packaged as AAV2/9 viruses, and produced with titers of 2.0E+13 particles per mL by OBio (Shanghai, China). Mice were anesthetized by 1% pentobarbital (50 mg/kg, i.p.) and mounted at stereotaxic apparatus (RWD68025). AAV-*TRE*-EYFP (2 μL) was injected into the corpus callosum (from bregma in mm, M-L: ±1.2, A-P: +0.5, D-V: -2.2) under the control of micropump (KDS310) at a speed of 0.05 μL/min. Injecting needles (Hamilton NDL ga33/30 mm/pst4) were withdrawn 10 min after injection. Infected brains were isolated 1 or 2 days later and brain slices were immunostained with anti-GFP antibody to enhance the visualization of the reporter protein.

### Electron Microscopy

Mice were anesthetized and transcardially perfused with 4% sucrose, 4% paraformaldehyde (PFA) and 2% glutaraldehyde in 0.1 M phosphate buffer (PB, pH 7.4). The brains and optic nerves were isolated carefully. The corpora callosa and prefrontal cortices were further dissected carefully under stereoscope. Tissues were post-fixed overnight at 4°C in 1% glutaraldehyde in 0.1 M PB. 24 h later, samples were washed by 0.1 M PB and osmicated with 2% osmium tetroxide 30-60 min at 4°C, washed by 0.1 M PB and by deionized H_2_O at 4°C, and dehydrated in graded (50-100%) ethanol. Samples were incubated with propylene oxide and embedded with embedding resins. Ultrathin sections were stained with 2% uranyl acetate at 4°C for 30 min, and then photographed with Tecnai 10 (FEI). EM images were analyzed by ImageJ (NIH). To eliminate the bias on circularity, *g*-ratio of each axon was calculated by the perimeter of axons (inner) divided by the perimeter of corresponding fibers (outer). Axonal diameters were normalized by perimeters through equation: diameter = perimeter/π. This procedure allows for inclusion of irregularly shaped axons and fibers and helps to eliminate biased measurement of diameters based on circularity. For quantitative analysis, cross sections of each neural tissue were divided into 5 areas, and more than two images, randomly selected from each area, were examined. The g-ratio of detectable myelinated axons in these images was measured. The numbers of detectable axons from each mouse were not the same. *g*-ratio from all detectable axons of one mouse was averaged and averaged *g*-ratio of three to five mice were used for statistical analysis.

### Immunofluorescence staining

Deeply anesthetized mice were transcardially perfused with 0.01 M PBS and then 4% PFA in 0.01 M PBS. Mouse brains were isolated and post-fixed in 4% PFA in 0.01 M PBS overnight at 4 °C, and then transferred into 20% and subsequently 30% sucrose in PBS overnight at 4 °C. Brains were then embedded in OCT (Thermo Fisher scientific) and sectioned into 20 μm on a cryostat sectioning machine (Thermo Fisher scientific, Microm HM525). Brain slices were incubated with a blocking buffer (10% fetal bovine serum and 0.2% Triton-X-100 in 0.01 M PBS) for 1 h at room temperature (RT), and then incubated at 4 °C overnight with primary antibodies diluted in the blocking buffer. The primary antibodies used were: GFP (1:500, Abcam, ab13970), CC1 (1:500, Abcam, ab16794), NG2 (1:200, Abcam, ab50009), Ki67 (1:400, Cell Signaling Technology, 9129), GFAP (1:2000, Millipore, MAB360), Iba1 (1:1000, Millipore, MABN92), MBP (1:100, Millipore, MAB382), TCF4 (1:500, Millipore, 05-511), Olig2 (1:500, Millipore, AB9610), TUJ1 (1:500, Sigma, T5076), RIP3 (1:500, QED, 2283), MLKL (1:500, Abgent, AP14272B). After washing three times with 0.1% Triton-X-100 in 0.01 M PBS, samples were incubated at RT for 1 h with Alexa-488 or -594 secondary antibody, and then washed and mounted on adhesion microscope slides (CITOTEST) with fluorescent mounting medium. Nuclear labeling was completed by incubating slices with DAPI (0.1 μg/mL, Roche) at RT for 5 min after incubation with secondary antibodies. Except for the antibody against NG2, antigen retrieval in 0.01 M sodium citrate buffer (pH 6.0) at 80-90 °C for 10 min was necessary before primary antibody incubation for brain slices to achieve definitive signals. Images were taken by a Zeiss LSM710 confocal microscope or a Nikon Eclipse 90i microscope. For cell counting based on immunostaining results, soma-shaped immunoreactive signals associated with a nucleus was counted as a cell. The immunostaining intensity was measured by ImageJ with background subtraction. Averaged intensity of areas with MBP signals was calculated as MBP intensity and normalized to that of controls. For MBP distribution analysis, images were applied with a threshold to eliminate the background, and the percentage of signal area in the total area was quantified as the signal distribution and normalized to that of controls.

### Luxol fast blue (LFB) staining

Brain slices were obtained as described in Immunofluorescence staining. After sufficient washing with 0.01 M PBS, the slices were transferred into a mixture of trichloromethane and ethanol (volume ratio 1:1) for 10 min and then 95% ethanol for 10 min. They were next incubated in 0.2% Luxol fast blue staining solution (0.2 g Solvent blue 38, 0.5 mL acetic acid, 95% ethanol to 100 mL) at 60 °C overnight. The next day, tissues were incubated for 5 min each in turn in 95% ethanol, 70% ethanol and ddH_2_O for rehydration, followed by incubation alternatively in 0.05% Li_2_CO_3_, 70% ethanol and ddH_2_O for differentiation until the contrast between the gray matter and white matter became obvious. After that, tissues were incubated for 10 min each in 95% and 100% ethanol to dehydrate, and then 5 min in dimethylbenzene to clear, before quickly mounting with neutral balsam mounting medium (CWBIO). All steps were operated in a ventilation cabinet. The LFB intensity in the corpus callosum was measured by Image J with background subtraction, and normalized to that of controls.

### TUNEL assay

Apoptotic cells were examined with terminal deoxynucleotidyl transferase (TdT)-mediated deoxyuridine triphosphate (dUTP) nick-end labeling (TUNEL) assay according to the manufacturer’s instructions (Vazyme; Yeasen). In brief, brain slices which were obtained as that described in Immunofluorescence staining were digested for 10 min by proteinase K (20 μg/mL) at RT. After washing twice with PBS, brain slices were incubated with Equilibration Buffer for 30 min at RT, and subsequently with Alexa Fluor 488-12-dUTP Labeling Mix for 60 min at 37°C. After washing with PBS three times, brain slices were stained with DAPI before being mounted under coverslips. For co-labeling of apoptotic nuclei in slices with immunofluorescence staining, TUNEL assay was performed after washing of the secondary antibody.

### Western blotting

Subcortical white matter tissues including the corpus callosum were isolated and homogenized. Homogenates in lysis buffer (10 mM Tris-Cl, pH 7.4, 1% NP-40, 0.5% Triton-X 100, 0.2% sodium deoxycholate, 150 mM NaCl, 20% glycerol, protease inhibitor cocktail) at a ratio of 2 mL per 100 mg tissue were lysed overnight in 4°C. Lysates were centrifuged at 12,000 g and 4°C for 15 min to get rid of the unsolved debris. Concentration of the supernatant was measured by BCA assay. Proteins in samples were separated by 6-12% SDS-PAGE, transferred to a Immobilon-P Transfer Membrane (Millipore), and then incubated with indicated primary antibodies diluted in blocking buffer at 4°C overnight after blocking by 5% non-fat milk solution in TBST (50 mM Tris, pH 7.4, 150 mM NaCl, 0.1% Tween 20) for 1 h at RT. The primary antibodies used were: pErbB3 (1:2500, Abcam, ab133459), pErbB4 (1:2500, Abcam, ab109273), pErbB2 (1:2500, Abgent, AP3781q), EGFR (1:5000, Epitomics, 1902-1), pEGFR (1:2500, Epitomics, 1727-1), GAPDH (1:5000, Huabio, EM1101), MBP (1:1000, Millipore, MAB382), ErbB3 (1:200, Santa Cruz Biotechnology, sc-285), ErbB4 (1:200, Santa Cruz Biotechnology, sc-283), ErbB2 (1:200, Santa Cruz Biotechnology, sc-284), Akt (1:5000, Cell signaling Technology, 9272), pAkt (1:3000, Cell signaling Technology, 9271), Erk1/2 (1:5000, Cell signaling Technology, 9102), pErk1/2 (1:5000, Cell signaling Technology, 4370), RIP3 (1:2000, QED, 2283), MLKL (1:2000, Abgent, AP14272B). For antibodies against phosphorylated proteins, 10% fetal bovine serum was used as a blocking buffer. Next day, the membranes were washed by TBST for three times and incubated with the secondary antibodies for 1 h at RT. Membranes were washed again and incubated with Immobilon Western Chemiluminescent HRPSubstrate (Millipore) for visualization of chemiluminescence by exposure to X-ray films or Bio-Rad GelDOCXR^+^ Imaging System. Intensities of protein bands were measured by ImageJ, and statistical analysis was performed after subtraction of the background intensity and normalization with controls in each batch of experiments.

### OPC cultures and differentiation *in vitro*

The cortices of mice at P2–P5 were isolated and sheared into pieces, and digested by 0.25% Trypsin in Ca2^+^Mg2^+^-free D-Hanks for 30 min at 37 °C. Trypsinized tissues were triturated and passed through 40 μm mesh. OPCs were obtained from the dissociated cells using MS Columns (Miltenyi Biotec 130-042-201) with magnetic beads (Miltenyi Biotec 130-101-547) to positively select PDGFRα^+^ cells according to the manufacturer’s recommendations. Isolated OPCs were plated into a poly-L-lysine-coated glass coverslips at a density of 20,000 cells per cm^2^ with DMEM/F12 supplemented with 10% fetal bovine serum. Four hours later, the medium was replaced with DMEM/F12 medium with 2% B27, 20 ng/mL bFGF and 10 ng/mL PDGFAA. For differentiation induction, the medium was replaced with DMEM/F12 medium with 2% B27 and 40 ng/mL triiodothyronine. For cells from *Sox*-ErbB2^V664E^ mice, 3 µg/mL Dox was added into the medium 24 h after plating.

### Statistical Analysis

All data were analyzed using Prism (Graphpad) and presented as mean ± s.e.m.. Unpaired two-tailed Student’s *t* test was used for analysis between two groups with one variable. Statistical significance was set at \**P* < 0.05, \*\**P* < 0.01, \*\*\**P* < 0.001. For western blotting and staining results, statistical analyses were performed after subtraction of the background intensity and normalization with controls in each batch of experiments to minimize the influences of batch-to-batch variations. Detailed statistical information for each experiment was included in the figure legends.

## Results

### ErbB overactivation caused inflammatory demyelination in *Plp*-ErbB2^V664E^ mice

To manipulate ErbB receptor activities specifically in oligodendrocytes *in vivo*, we employed tetracycline-controlled systems whose induction or blockade depends on the presence of doxycycline (Dox). We generated *Plp*-tTA;*TRE*-ErbB2^V664E^ (*Plp*-ErbB2^V664E^) bi-transgenic mice by crossing *Plp*-tTA with *TRE*-ErbB2^V664E^ mice. Breeding pairs, as well as their offspring, were fed with Dox to ‘Tet-off’ the *TRE*-controlled transcription. Dox was withdrawn when the offspring were weaned on P21 (Fig. 1*A*). Among ErbB1-4 receptors, ErbB2 that does not bind to any known ligand is the preferred partner to other ligand-bound ErbB members. ErbB2^V664E^ contains an amino acid mutation (Vla_664_/Glu_664_) within the transmembrane domain facilitating its dimerization with other ErbB receptors and potentiating their downstream signaling (Chen et al., 2017). ErbB2^V664E^ expression was significantly induced in the white matter of *Plp*-ErbB2^V664E^ mice within 1 day post Dox-withdrawal (dpd) (Fig. 1*B*). Notably, *Plp*-ErbB2^V664E^ mice around P35 exhibited severe ataxia while walking on a grid panel (Fig. 1*C*). Moreover, *Plp*-ErbB2^V664E^ mice showed difficulty in rolling over, indicating severely impaired motor coordination. This was unexpected as this phenotype has not been reported for NRG1- or hEGFR- overexpressing mice (Aguirre et al., 2007; Brinkmann et al., 2008).

**Figure 1.**
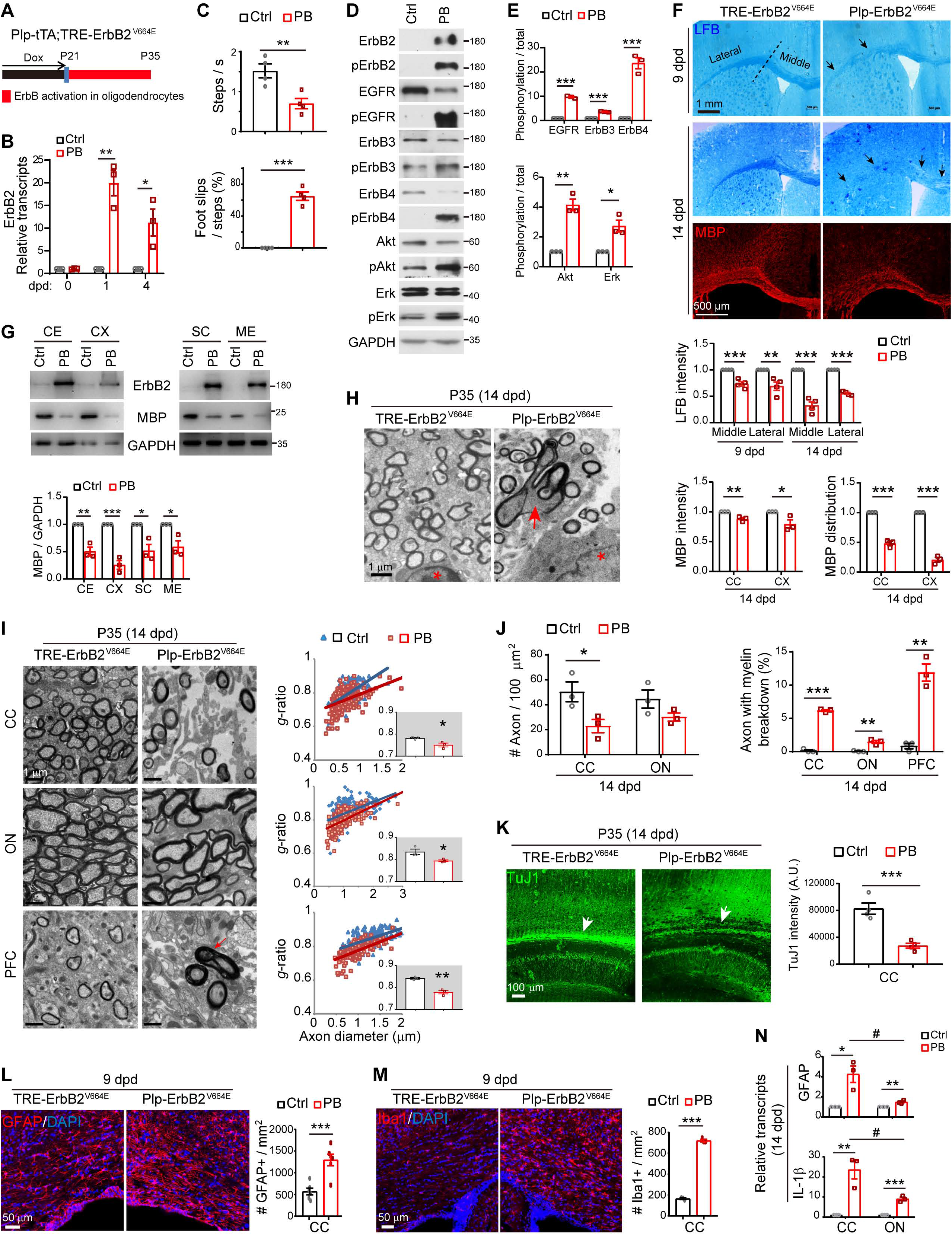
ErbB overactivation in *Plp*-ErbB2^V664E^ mice induced inflammatory demyelination. Unless otherwise indicated, quantitative data were presented in the graphs as mean ± s.e.m., and analyzed by unpaired *t* test. **A,** Dox treatment setting for indicated mice and littermate controls. **B,** Real-time RT-PCR results of ErbB2 expression at indicated day post Dox withdrawal (dpd). Statistical information for 1 dpd, *t*_(4)_ = 6.72, *P* = 0.0025; for 4 dpd, *t*_(4)_ = 3.432, *P* = 0.026. **C,** Walking speed and percentage of foot slips of *Plp*-ErbB2^V664E^ mice and littermate controls at P35 with 14 dpd in the grid walking test. n = 4 mice for each group. For steps per second, *t*_(6)_ = 3.773, *P* = 0.0093; for foot slip percentage, *t*_(6)_ = 12.31, *P* < 0.0001. **D**, Western blotting of indicated proteins in white matter isolated from *Plp*-ErbB2^V664E^ (PB) mice in comparison with that from littermate control mice (Ctrl). Activation status of each ErbB receptor or downstream signaling protein was examined by western blotting with the specific antibody against its phosphorylated form. **E,** Quantitative data of western blotting results. For EGFR, *t*_(4)_ = 27.64, *P* < 0.0001; for ErbB3, *t*_(4)_ = 19.98, *P* < 0.0001; for ErbB4, *t*_(4)_ = 10.06, *P* = 0.00055; for Akt, *t*_(4)_ = 8.096, *P* = 0.0013; for Erk, *t*_(4)_ = 4.353, *P* = 0.012. **F,** LFB staining and MBP immunostaining results of coronal sections through the genu of the corpus callosum in *Plp*-ErbB2^V664E^ and control mice. Black arrows indicate the lower staining intensity of myelin stained in the corpus callosum. Statistical information for quantitative data of LFB intensity: the middle part at 9 dpd, *t*_(6)_ = 6.345, *P* = 0.00072; the lateral part at 9 dpd, *t*_(6)_ = 3.914, *P* = 0.0078; the middle part at 14 dpd, *t*_(6)_ = 9.89, *P* < 0.0001; the lateral part at 14 dpd, *t*_(6)_ = 23.07, *P* < 0.0001. Statistical information for MBP intensity: Corpus callosum (CC), *t*_(4)_ = 4.884, *P* = 0.0081; Cortex (CX), *t*_(4)_ = 2.834, *P* = 0.047. Statistical information for MBP distribution: CC, *t*_(4)_ = 14.53, *P* = 0.00013; CX, *t*_(4)_ = 20.51, *P* < 0.0001. **G,** Western blotting results of MBP and ErbB2 in multiple brain regions (CX, cortex; CE, cerebellum; SC, spinal cord; ME, medulla) isolated from *Plp*-ErbB2^V664E^ (PB) and littermate control mice (Ctrl) at 14 dpd. GAPDH served as a loading control. Statistical information for CE, *t*_(4)_ = 6.35, *P* = 0.0032; for CX, *t*_(4)_ = 9.243, *P* = 0.00076; for SC, *t*_(4)_ = 4.118, *P* = 0.0146; for ME, *t*_(4)_ = 3.634, *P* = 0.022. **H**, Representative EM images showed that myelin sheath ruptured and broke down in *Plp-*ErbB2^V664E^ mice at 14 dpd (red arrow). Note the associated nuclei (red asterisk) showed no chromatin condensation and nucleation. **I,** EM images of the corpus callosum (CC), optic nerve (ON), and prefrontal cortex (PFC) from *Plp*-ErbB2^V664E^ and littermate controls at 14 dpd. Quantitative data were shown for *g*-ratio analysis of myelinated axons detected by EM. Averaged *g*-ratio for each mouse were plotted as insets, presented as mean ± s.e.m., and analyzed by unpaired *t* test. For CC, *t*_(4)_ = 3.412, *P* = 0.027; for ON, *t*_(4)_ = 3.083, *P* = 0.037; for PFC, *t*_(4)_ = 7.11, *P* = 0.0021. The red arrow indicates the axon with myelin breakdown. **J,** The densities of myelinated axons, as well as the percentages of axons with myelin breakdown, examined by EM in different brain regions of *Plp*-ErbB2^V664E^ (PB) and littermate control mice (Ctrl) at 14 dpd were quantified. Statistical information for myelinated-axon density, in CC, *t*_(4)_ = 2.683, *P* = 0.046; in ON, *t*_(4)_ = 1.818, *P* = 0.143. For the percentage of axons with myelin breakdown, in CC, *t*_(4)_ = 29.32, *P* < 0.0001; in ON, *t*_(4)_ = 6.108, *P* = 0.0036; in PFC, *t*_(4)_ = 8.125, *P* = 0.0012. **K,** Axons were reduced in the subcortical white matter (white arrows) of *Plp*-ErbB2^V664E^ mice at 14 dpd. Sagittal sections of *Plp*-ErbB2^V664E^ and littermate control mice were immunostained by monoclonal antibody TuJ1. Subcortical staining intensities were measured and plotted as mean ± s.e.m., and analyzed by unpaired *t* test. *t*_(4)_ = 6.019, *P* = 0.0009. **L, M,** Astrocytes (GFAP^+^) and microglia (Iba1^+^) examined in the subcortical white matter of indicated mice by immunostaining. Cell densities in the corpus callosum were quantified, and data were presented as mean ± s.e.m., and analyzed by unpaired *t* test. In **L**, *t*_(10)_ = 4.753, *P* = 0.0008. In **M**, *t*_(4)_ = 36.4, *P* < 0.0001. **N,** Real-time RT-PCR results of GFAP and IL-1β transcripts in the corpus callosum (CC) and optic nerve (ON). *, *P* < 0.05; **, *P* < 0.01; ***, *P* < 0.001, PB *vs* Ctrl. #, *P* < 0.05, CC *vs* ON. See also Extended data Figure 1-1 and Figure 1-2.

In *Plp*-ErbB2^V664E^ mice at P21 before Dox withdrawal, activities of endogenous ErbB receptors and the downstream Akt and Erk (MAPK) in white matter were not altered (Extended data Fig. 1-1*A*). In *Plp*-ErbB2^V664E^ mice at P35 with 14 dpd, with the expression and phosphorylation of ectopic ErbB2^V664E^, endogenous ErbB receptors in the white matter, including both EGFR that mediates EGF signaling and ErbB3/ErbB4 that mediate NRG signaling, were strikingly phosphorylated and activated (Fig. 1*D, E*). Activating ErbB receptors stimulated downstream signaling including both Erk and Akt, as their phosphorylation significantly increased in the white matter of *Plp*-ErbB2^V664E^ mice (Fig. 1*D*, E).

Overactivation of ErbB receptors caused lower myelin staining intensity as exhibited in the corpus callosum of *Plp*-ErbB2^V664E^ mice at 9 dpd after Luxol fast blue (LFB) staining (Fig. 1*F*). White matter tracts in the lateral part of corpus callosum of *Plp*-ErbB2^V664E^ mice at 9 dpd were obviously fragmented (black arrows). Notably, at 14 dpd, myelin loss became more evident throughout the corpus callosum, as that LFB staining intensities dropped dramatically in both the middle and lateral parts (Fig. 1*F*). Immunostaining of myelin basic protein (MBP) also revealed dramatic myelin loss in the corpus callosum at 14 dpd (Fig. 1*F*), suggesting that *Plp*-ErbB2^V664E^ mice were undergoing CNS demyelination after Dox withdrawal. Western blotting revealed that the induced ErbB2^V664E^ expression was associated with MBP loss in multiple brain regions of *Plp*-ErbB2^V664E^ mice (Fig. 1*G*), indicating a global CNS demyelination. Indeed, the electron microscopic (EM) examination revealed that myelin sheaths of some axons in *Plp*-ErbB2^V664E^ mice ruptured or underwent breakdown (Fig. 1*H*, *I*, *J*). Due to demyelination, only a few intact axons were detected in the midline of the corpus callosum of *Plp*-ErbB2^V664E^ mice at 14 dpd (Fig. 1*I*, J). When axonal tracts in the corpus callosum of *Plp*-ErbB2^V664E^ mice at 14 dpd were immunostained by TuJ1, the antibody recognizing neuronal specific β-tubulin III, the immunoreactivity dramatically reduced (Fig. 1*K*). Inflammatory demyelination is usually complicated and aggravated by the pathological responses from nearby astrocytes and microglia (Bradl and Lassmann, 2010; Burda and Sofroniew, 2014). Indeed, in the white matter of *Plp*-ErbB2^V664E^ mice, astrogliosis and microgliosis were revealed after Dox withdrawal (Extended data Fig. 1-1*B*, *C*; Fig. 1*L*, *M*). It was likely that inflammation exacerbated myelin degradation and axon degeneration in the corpus callosum, given the fact that most axons were preserved in the optic nerves where inflammatory status, as indicated by transcription of GFAP and pro-inflammatory cytokine IL-1β, was significantly milder than that in the corpus callosum of the same mice at 14 dpd (Fig. 1*N*).

### *Plp*-ErbB2^V664E^ white matter exhibited hypermyelination prior to demyelination

Interestingly, despite the demyelination, the detectable axons, throughout the corpus callosum, optic nerve, and prefrontal cortex in *Plp*-ErbB2^V664E^ mice, were hypermyelinated (Fig. 1*I*). Myelinated axons detected in the brain of *Plp*-ErbB2^V664E^ mice had significantly smaller *g*-ratio (axon diameter/fiber diameter), a quantitative indication of myelin thickness for individual axons with different diameters (Fig. 1*I*). Myelin thickness showed no difference between *TRE*-ErbB2^V664E^ and littermate *Plp*-tTA, two single transgenic control mice, after Dox withdrawal (Extended data Fig. 1-2*A*). Therefore, hypermyelination of detectable axons in *Plp*-ErbB2^V664E^ mice was a result of ErbB signaling overactivation.

Hypermyelination of individual axons in *Plp*-ErbB2^V664E^ mice phenocopied that observed in NRG1-overexpressing mice (Brinkmann et al., 2008), as well as that in the transgenic mice with Akt or Erk signaling overactivation in oligodendrocytes (Harrington et al., 2010; Ishii et al., 2013). We immunostained MBP in *Plp*-ErbB2^V664E^ mice, and found that myelin structures and distributions remained mostly normal in the middle part of corpus callosum at 9 dpd (Fig. 2*A*, inset 2’ *vs* inset 2), despite that MBP loss and myelin fragmentation were prominent at the cortical projection tips in the same mice (Fig. 2*A*, *C*). An examination of the ultrastructure in these mice by EM revealed similar results that most axons were intact in the midline of the corpus callosum, although they have been significantly hypermyelinated (Fig. 2*B*, *D*). These results confirmed that hypermyelination occurred early in *Plp*-ErbB2^V664E^ mice, consistent with the reported function of ErbB signaling overactivation (Brinkmann et al., 2008; Harrington et al., 2010; Ishii et al., 2013). Overactivation of ErbB receptors activates multiple downstream signaling, and transduce more active intracellular signaling than transduction through the increase of ligands. Rapid demyelination and axonal degeneration after 9 dpd may be attributed to inflammation that exacerbated the pathological progression in *Plp*-ErbB2^V664E^ mice (Burda and Sofroniew, 2014).

**Figure 2.**
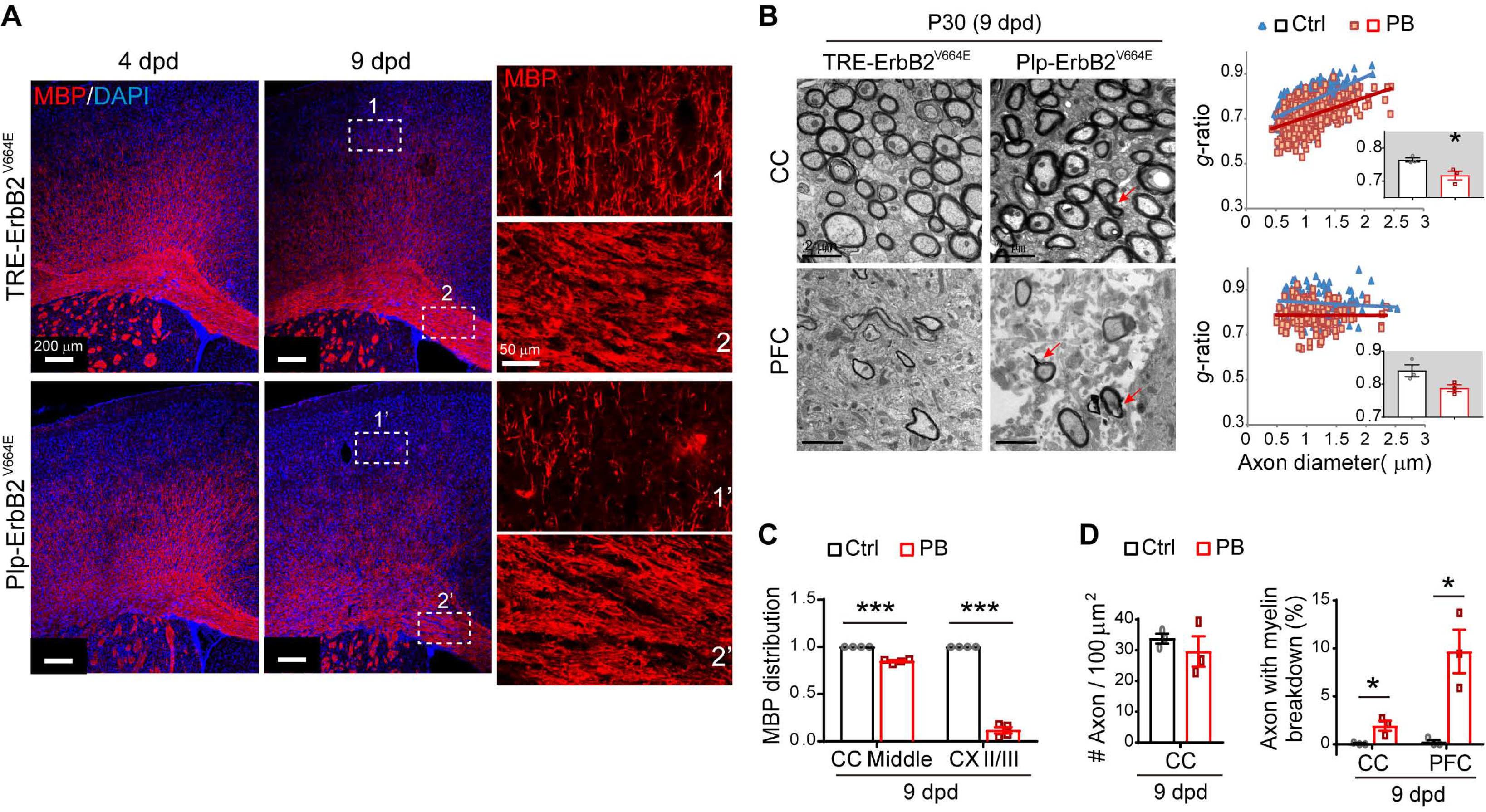
*Plp*-ErbB2^V664E^ white matter exhibited hypermyelination prior to demyelination. **A**, MBP immunostaining of brain slices from indicated mice. Note the myelin structure and distribution exhibited relatively more normal in the middle part of corpus callosum (inset 2’) than that in the layer II/III of cortex (inset 1’), in *Plp*-ErbB2^V664E^ mice at 9 dpd. **B,** EM examination of axons in the midline of the corpus callosum (CC) and the prefrontal cortex (PFC) in *Plp*-ErbB2^V664E^ mice at 9 dpd. *g*-ratio was calculated for myelinated axons and averaged *g*-ratio were analyzed by unpaired *t* test (inset). For CC, *t*_(4)_ = 3.226, *P* = 0.032; for PFC, *t*_(4)_ = 2.529, *P* = 0.065. Red arrows indicate the axons with myelin breakdown. **C,** Quantitative data of MBP signal distribution in immunostaining results. Data were presented as mean ± s.e.m., and analyzed by unpaired *t* test. In middle CC, *t*_(6)_ = 13.11, *P* < 0.0001; in cortex layer II/III, *t*_(6)_ = 26.31, *P* < 0.0001. **D,** The densities of myelinated axons as well as the percentages of axons with myelin breakdown in EM analysis. Data were presented as mean ± s.e.m., and analyzed by unpaired *t* test. For myelinated-axon density in CC, *t*_(4)_ =0.805, *P* = 0.466. For axons with myelin breakdown at 9 dpd examined by EM: in CC, *t*_(4)_ = 3.567, *P* = 0.023; in PFC, *t*_(4)_ = 4.146, *P* = 0.014.

### ErbB overactivation induced oligodendrocyte degeneration but OPC regeneration in *Plp*-ErbB2^V664E^ mice

To investigate the pathological mechanism, we examined the post-mitotic oligodendrocytes by immunostaining with antibody CC1, and found a decreasing number of intact CC1^+^ cells in the corpus callosum of *Plp*-ErbB2^V664E^ mice starting from 6 dpd (Fig. 3*A*, *C*). Meanwhile, OPC (NG2^+^Olig2^+^) numbers were dramatically increased (Fig. 3*B*, *C*). These disproportionate oligodendrocyte lineage state changes were consistent with the previous reports that demyelination induces OPC regeneration (Aguirre et al., 2007; Bradl and Lassmann, 2010). We examined OPC proliferation by co-immunostaining for Olig2 and Ki67, the proliferation marker. In normal condition after early development, most proliferative OPCs (Ki67^+^Olig2^+^) resided in the subventricular zone, with a few in the trunk of corpus callosum (Fig. 3*D*). In *Plp*-ErbB2^V664E^ mice starting from 6 dpd, dramatic proliferation of OPCs was observed within the trunk of corpus callosum instead of the subventricular zone (Fig. 3*D*, *E*), further indicating the pathogenetic state of myelin. The non-specifically stained hemorrhagic spots (white asterisk) in the cortex of *Plp*-ErbB2^V664E^ mice also indicated the inflammatory pathological status. The pathological events, including myelin (MBP^+^) breakdown, oligodendrocytes (CC1^+^) degeneration, and OPC (PDGFRα^+^) regeneration, were also observed in *Plp*-ErbB2^V664E^ mice at P70 with 10 dpd (Extended data Fig. 3-1*A-E*). These pathological events were correlated with inflammation as well (Extended data Fig. 3-1*F-H*), despite that the inflammatory profiles (GFAP^+^ cell numbers reduced while Iba1^+^ cell numbers increased significantly in the corpus callosum) were not the same as those in mice at P30 with 9 dpd (both GFAP^+^ and Iba1^+^ cell numbers increased significantly). There were even more hemorrhagic spots (white asterisk) in the cortex of adult *Plp*-ErbB2^V664E^ mice, indicating that the adult brains were more vulnerable to inflammation. Therefore, ErbB overactivation induced by *Plp*-tTA was detrimental to oligodendrocytes, causing inflammatory demyelination at both adolescence and adulthood.

**Figure 3.**
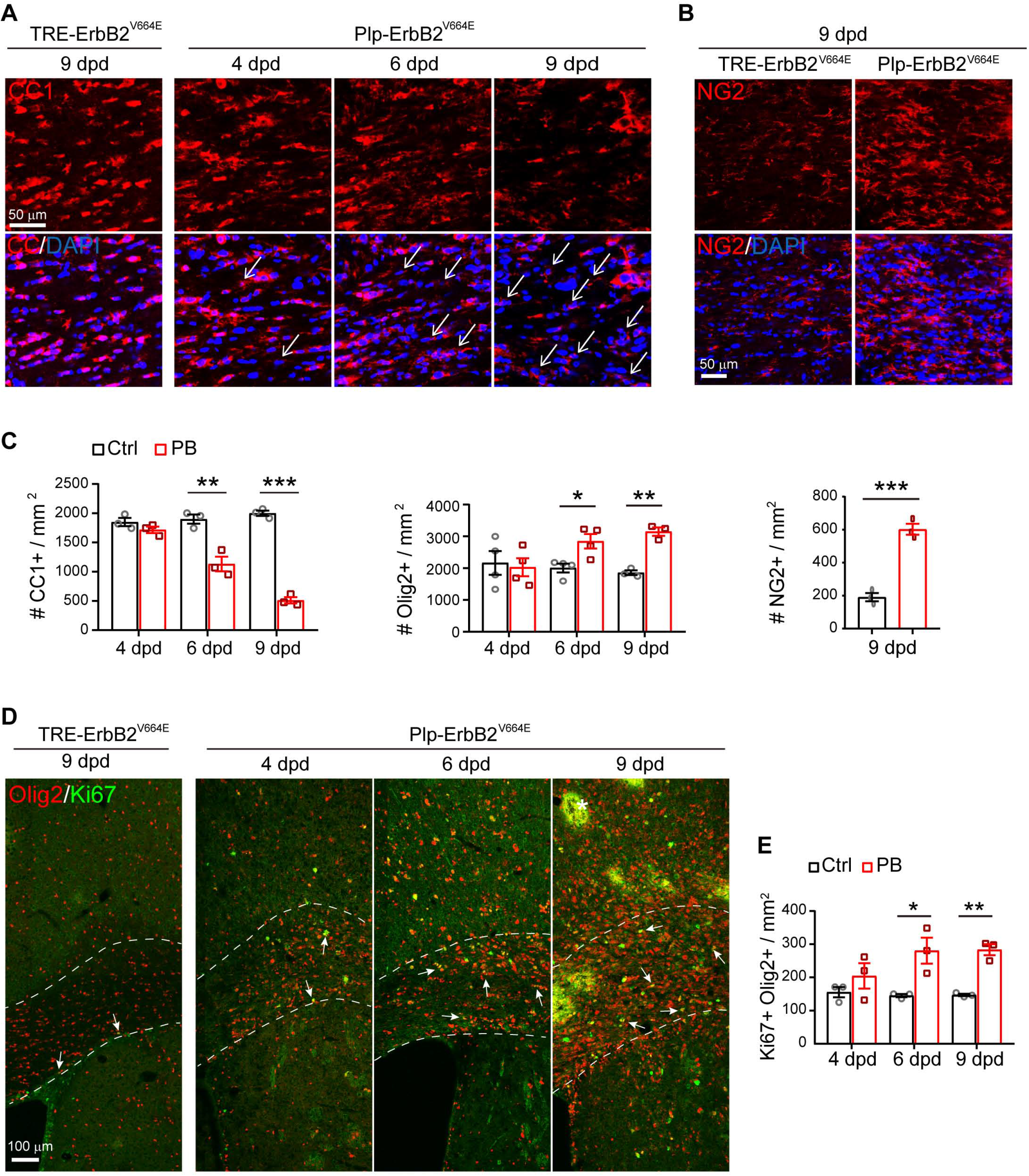
ErbB overactivation induced oligodendrocyte degeneration but OPC regeneration in *Plp*-ErbB2^V664E^ mice. **A,** The numbers of degenerating oligodendrocytes (represented by nuclei associated with CC1^+^ cell debris, white arrows) increased in the corpus callosum of *Plp*-ErbB2^V664E^ mice starting from 6 dpd as revealed by CC1 immunostaining. The densities of remained intact CC1^+^ cells were reduced. **B,** Immunostaining results of NG2 in the corpus callosum of indicated mice at 9 dpd. **C,** Quantitative data of immunostaining results in *Plp*-ErbB2^V664E^ (PB) and control mice (Ctrl) with indicated Dox treatments were presented as mean ± s.e.m., and analyzed by unpaired *t* test. For CC1^+^ density: at 4 dpd, *t*_(4)_ = 1.485, *P* = 0.212; for 6 dpd, *t*_(4)_ = 5.203, *P* = 0.0065; for 9 dpd, *t*_(4)_ = 20.95, *P* < 0.0001. For Olig2^+^ density: at 4 dpd, *t*_(6)_ = 0.2923, *P* = 0.780; at 6 dpd, *t*_(6)_ = 3.16, *P* = 0.0196; at 9 dpd, *t*_(4)_ = 8.563, *P* = 0.001. For NG2^+^ density at 9 dpd, *t*_(4)_ = 9.912, *P* = 0.0006. **D,** Immunostaining results of Olig2 with Ki67 in the corpus callosum of indicated mice. Note there were non-specifically stained hemorrhagic spots (white asterisk), which is the consequence of inflammation, in the brain slices from *Plp*-ErbB2^V664E^ mice at 9 dpd. **E,** Quantitative data of Ki67^+^Olig2^+^ cell density in *Plp*-ErbB2^V664E^ (PB) and control mice (Ctrl) with indicated Dox treatments were present as mean ± s.e.m., and analyzed by unpaired *t* test. At 4 dpd, *t*_(4)_ = 1.187, *P* = 0.301; at 6 dpd, *t*_(4)_ = 3.428, *P* = 0.027; at 9 dpd, *t*_(4)_ = 8, *P* = 0.0013. See also Extended data Figure 3-1.

### ErbB overactivation caused oligodendrocyte number reduction and non-inflammatory hypomyelination in *Sox10*-ErbB2^V664E^ mice

Next, we examined the effects of ErbB overactivation on CNS myelin by a ‘Tet-on’ system generated in *Sox10*^+/rtTA^;*TRE*-ErbB2^V664E^ (*Sox10*-ErbB2^V664E^) mice (Fig. 4*A*). ErbB2^V664E^ expression was also significantly induced in the white matter within 1 day with Dox-feeding (dwd), suggesting similarly prompt responses of tTA/rtTA transcription in *Sox10*-ErbB2^V664E^ and *Plp*-ErbB2^V664E^ mice (Fig. 4*B*). *Sox10*-ErbB2^V664E^ mice with Dox feeding from P21 developed severe motor dysfunction, including ataxia and tremors, and died around P35. As a result, *Sox10*-ErbB2^V664E^ and littermate control mice were investigated at P30 with 9 dwd. These mice had smaller body sizes at P30 and walked with difficulty on a grid panel (Fig. 4*C*). Interestingly, *Sox10*-ErbB2^V664E^ mice performed normal rolling over. Despite walking slowly in the grid walking test, *Sox10*-ErbB2^V664E^ mice did not exhibit as many foot slips as *Plp*-ErbB2^V664E^ mice did. On the other hand, *Plp*-ErbB2^V664E^ mice did not exhibit tremors. The tremors in *Sox10*-ErbB2^V664E^ mice should be ascribed to the hypomyelination of peripheral nerves (Extended data Fig. 4-1*A-H*).

**Figure 4.**
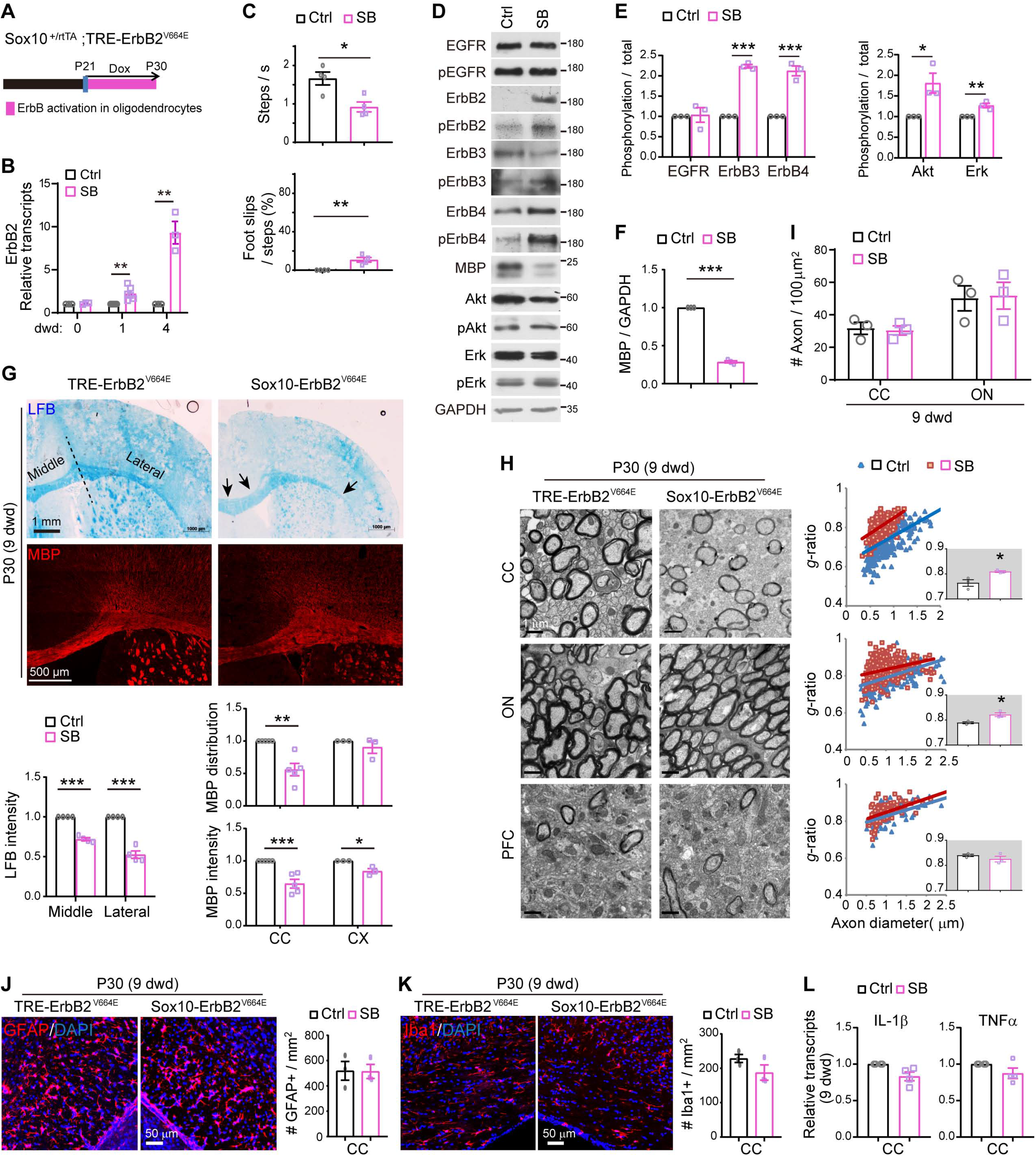
ErbB overactivation caused non-inflammatory hypomyelination in *Sox10*-ErbB2^V664E^ mice. Unless otherwise indicated, quantitative data were presented as mean ± s.e.m., and analyzed by unpaired *t* test. **A,** Dox treatment setting for indicated mice and littermate controls. **B,** Real-time RT-PCR results of ErbB2 expression at indicated day with Dox treatment (dwd). Statistical information for 1 dpd, *t*_(10)_ = 4.081, *P* = 0.0022; for 4 dpd, *t*_(4)_ = 6.37, *P* = 0.0031. **C,** Walking speed and percentage of foot slips of *Sox10*-ErbB2^V664E^ mice and littermate controls at 9 dwd in the grid walking test. n = 4 mice for each group. Statistical information for steps per second, *t*_(6)_ = 3.504, *P* = 0.0128; for foot slip percentage, *t*_(6)_ = 4.429, *P* = 0.0044. **D,** Western blotting of indicated proteins in white matter tissues isolated from *Sox10*-ErbB2^V664E^ (SB) mice in comparison with that from littermate control mice (Ctrl). Activation status of each ErbB receptor or downstream signaling protein was examined by western blotting with the specific antibody against its phosphorylated form. **E, F,** Quantitative data of western blotting results. In **E**, statistical information for EGFR, *t*_(4)_ = 0.1983, *P* = 0.852; for ErbB3, *t*_(4)_ = 28.34, *P* < 0.0001; for ErbB4, *t*_(4)_ = 9.181, *P* = 0.00078; for Akt, *t*_(4)_ = 3.380, *P* = 0.028; for Erk, *t*_(4)_ = 4.899, *P* = 0.008. In **F**, for MBP, *t*_(4)_ = 48.82, *P* < 0.0001. **G**, LFB staining and MBP immunostaining results of coronal sections through the genu of the corpus callosum in *Sox10*-ErbB2^V664E^ (SB) and control mice (Ctrl). Black arrows indicate the lower staining intensity of myelin stained in the corpus callosum. Statistical information for quantitative data of LFB intensity: the middle part, *t*_(6)_ = 15.17, *P* < 0.0001; the lateral part, *t*_(6)_ = 10.23, *P* < 0.0001. Statistical information for MBP intensity: CC, *t*_(8)_ = 5.14, *P* = 0.0009; CX, *t*_(4)_ = 4.091, *P* = 0.015. Statistical information for MBP distribution: CC, *t*_(8)_ = 4.622, *P* = 0.0017; CX, *t*_(4)_ = 0.997, *P* = 0.375. **H,** EM images of the corpus callosum (CC), optic nerve (ON), and prefrontal cortex (PFC) from *Sox10*-ErbB2^V664E^ and littermate controls at 9 dwd. Quantitative data were shown for *g*-ratio analysis of myelinated axons detected by EM. Averaged *g*-ratio for each mouse were plotted as insets, presented as mean ± s.e.m., and analyzed by unpaired *t* test. For CC, *t*_(4)_ = 3.295, *P* = 0.0301; for ON, *t*_(4)_ = 3.775, *P* = 0.0195; for PFC, *t*_(4)_ = 1.196, *P* = 0.298. **I,** The densities of myelinated axons examined by EM in different brain regions of *Sox10*-ErbB2^V664E^ (SB) and littermate control mice (Ctrl) at 9 dwd were quantified. Statistical information for CC, *t*_(4)_ = 0.2773, *P* = 0.795; for ON, *t*_(4)_ = 0.1455, *P* = 0.891. **J, K,** Astrocytes (GFAP^+^) and microglia (Iba1^+^) examined in the subcortical white matter of indicated mice by immunostaining. Cell densities in the corpus callosum were quantified, and data were presented as mean ± s.e.m., and analyzed by unpaired *t* test. In **J**, *t*_(4)_ = 0.0501, *P* = 0.962. In **K**, *t*_(4)_ = 1.637, *P* = 0.178. **L**, Real-time RT-PCR results of IL-1β and TNFα transcripts in the corpus callosum (CC). Data were presented as mean ± s.e.m., and statistical analysis by unpaired *t* test revealed no differences. See also Extended data Figure 4-1, Figure 1-1, and Figure 1-2.

On P21 before Dox treatment, no ErbB2^V664E^ was detected in the white matter of *Sox10*-ErbB2^V664E^ mice, indicating there was no leaky expression of the transgene. Consistently, there were similar protein and phosphorylation levels of endogenous ErbB receptors as well as downstream Akt and Erk (Extended data Fig. 1-1*D*). After 9 dwd, western blotting revealed the expression and phosphorylation of ectopic ErbB2^V664E^ accompanied with increases in phosphorylation of ErbB3 and ErbB4, but not that of EGFR, in the white matter of *Sox10*-ErbB2^V664E^ mice (Fig. 4*D*, *E*). Even though only the activities of ErbB3 and ErbB4, the receptors to the NRG family ligands, were increased in the white matter of *Sox10*-ErbB2^V664E^ mice, both downstream Erk and Akt signaling were activated (Fig. 4*D*, *E*). With the ErbB signaling overactivation, brain slices stained by LFB or immunostained by MBP antibody exhibited lower staining intensity in the corpus callosum of *Sox10*-ErbB2^V664E^ mice (Fig. 4*G*), consistent with the lower MBP levels detected by western blotting (Fig. 4*D*, *F*). Unexpectedly, the examination of the ultrastructure of *Sox10*-ErbB2^V664E^ white matter by EM did not find hypermyelination or myelin breakdown, but revealed that the axons in the corpus callosum and optic nerve exhibited thinner myelin with significantly increased *g*-ratio (Fig. 4*H*). *Sox10*^+/rtTA^ is a knock-in mouse line, so that the allele with *Sox10*-rtTA would not transcribe *Sox10* mRNA (Ludwig et al., 2004). We analyzed the ultrastructure of myelinated axons in two control mice, *Sox10*^+/rtTA^ and littermate *TRE*-ErbB2^V664E^ mice, at P30 with 9 dwd and did not observe any differences (Extended data Fig. 1-2*B*). Therefore, we can exclude the possible effect of haploinsufficiency of Sox10 on late postnatal myelin development.

It is notable that in *Sox10*-ErbB2^V664E^ mice, the numbers of myelinated axons were not altered (Fig. 4*I*), and myelin sheaths exhibited normal morphology (Fig. 4*H*). Moreover, neither microgliosis nor astrogliosis was detected (Extended data Fig. 1-1*E*, *F*; Fig. 4*J*, *K*). Consistently, there was no increases in pro-inflammatory cytokine (IL-1β and TNFα) expression (Fig. 4*L*). Because there was no indication of inflammatory pathogenesis, we can conclude that thinner myelin in *Sox10*-ErbB2^V664E^ white matter was caused by developmental deficits not pathological conditions. Therefore, ErbB overactivation in *Sox10*-ErbB2^V664E^ mice induced CNS hypomyelination rather than inflammatory demyelination.

We further examined the states of oligodendrocytes and found the numbers of Olig2^+^, NG2^+^, and CC1^+^ cells all significantly decreased in the corpus callosum of *Sox10*-ErbB2^V664E^ mice with 6 and more dwd (Fig. 5*A*-*C*). The number reduction of NG2^+^ and CC1^+^ cells was proportional to each other and to that of Olig2^+^ cells, confirming that the deficits in *Sox10*-ErbB2^V664E^ mice were development-dependent. OPC proliferation decreased in the corpus callosum (Fig. 5*D*), further supporting that the oligodendrocyte lineage states in *Sox10*-ErbB2^V664E^ white matter were different from that in *Plp*-ErbB2^V664E^ white matter. As neither OPC nor oligodendrocyte differences were observed between littermate *Sox10*^+/rtTA^ and *TRE*-ErbB2^V664E^, two control mice, at P30 with 9 dwd (Extended data Fig. 5-1*A*,*B*), the developmental deficits in *Sox10*-ErbB2^V664E^ mice should be attributed to ErbB overactivation in oligodendrocytes.

**Figure 5.**
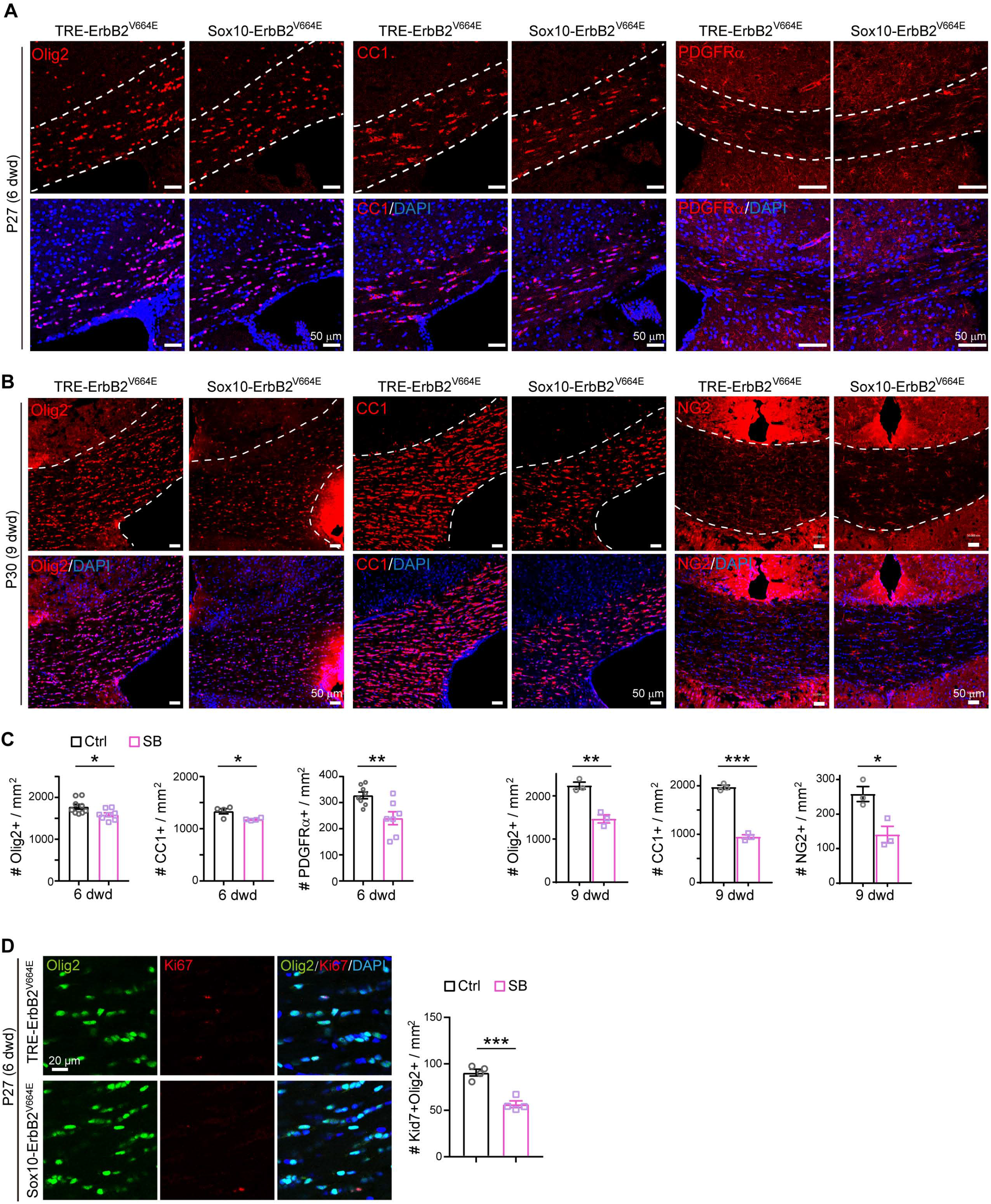
ErbB overactivation induced oligodendrocyte number reduction in *Sox10*-ErbB2^V664E^ mice. **A, B,** Olig2^+^, CC1^+^, and NG2^+^ cells in the corpus callosum of indicated mice at P27 with 6 dwd (**A**) or at P30 with 9 dwd (**B**) were examined by immunostaining. **C,** Statistic results of Olig2^+^, CC1^+^, and NG2^+^ cell densities in the corpus callosum of *Sox10*-ErbB2^V664E^ (SB) mice and littermate controls (Ctrl) with 6 dwd or 9 dwd. Data were from repeated immunostaining of 3 mice for each group, presented as mean ± s.e.m., and analyzed by unpaired *t* test. Statistical information for 6 dwd: Olig2^+^, *t*_(15)_ = 2.543, *P* = 0.0225; CC1^+^, *t*_(6)_ = 3.006, *P* = 0.0238; PDGFRα^+^, *t*_(13)_ = 3.236, *P* = 0.0065. For 9 dwd: Olig2^+^, *t*_(4)_ = 6.236, *P* = 0.0034; CC1^+^, *t*_(4)_ = 16.92, *P* < 0.0001; NG2^+^, *t*_(4)_ = 3.634, *P* = 0.0221. **D,** Immunostaining results of Olig2 with Ki67 in the corpus callosum of *Sox10*-ErbB2^V664E^ (SB) and control mice (Ctrl) at P27 with 6 dwd. Statistical information for quantitative data of Ki67^+^Olig2^+^ cell density: *t*_(6)_ = 6.629, *P* = 0.0006. See also Extended data Figure 5-1.

### *Plp*-tTA targeted mainly MOs whereas *Sox10*^+/rtTA^ targeted OPC-NFOs

To understand the cellular mechanisms of inflammatory demyelination (*Plp*-ErbB2^V664E^ mice) and non-inflammatory hypomyelination (*Sox10*-ErbB2^V664E^ mice) induced by ErbB overactivation, we had to first determine the cellular targeting preferences of Dox-dependent transgenes in these two mouse models. The finding that *Plp*-ErbB2^V664E^ and *Sox10*-ErbB2^V664E^ mice had few overlapping phenotypes was unexpected since that Sox10 is reported to express throughout the oligodendrocyte lineage, and *Sox10*^+/rtTA^ knock-in mice have been used to investigate all oligodendrocyte lineage cells (Wegener et al., 2015). On the other hand, *Plp*-tTA transgenic mice have been reported to target MOs in the adult brain (Inamura et al., 2012); however, *Plp* gene transcribes a shorter splicing isoform DM20 in OPCs and NFOs during embryonic and early postnatal development (Trapp et al., 1997). The previous report for *Sox10*^+/rtTA^-targeting specificity was concluded by using GFP reporter mice with Dox treatment for several weeks (Wegener et al., 2015). GFP is highly stable with a ∼26 h half-life (Corish and Tyler-Smith, 1999). In contrast, transactivator tTA has a half-life less than 6.5 h (Chassin et al., 2019). Therefore, GFP reporter generated in the original tTA/rtTA expressing cells can accumulate into the following cellular stages when tTA/rtTA has degraded, and thus the reporter-containing cells consist of tTA/rtTA-expressing cells and their progeny. In oligodendrocyte lineage, terminal differentiated OPCs can differentiate into NFOs in as quickly as 2.5 h and NFOs can mature in 1 day (Xiao et al., 2016). Because the induction of the ‘Tet-on’ or ‘Tet-off’ system by Dox feeding or Dox withdrawal has a delayed effect on gene expression, to examine the cellular targeting preferences under a condition with limited derivation from original tTA/rtTA-expressing cells, we acutely delivered a *TRE*-controlled fluorescence reporter carried by an adeno-associated virus (AAV) into the mouse brains at P14 or P35. In these AAV-*TRE*-YFP-infecting mice, rtTA- or tTA-targeting cells were pulse-labeled, allowing us to trace their differentiation states within a short time window.

*Plp*-tTA mice were raised with no Dox feeding, whereas *Sox10*^+/rtTA^ were fed with Dox for 3 days before the stereotaxic injection (Fig. 6*A*). One or 2 days after virus injection, the reporter-containing (YFP^+^) cells were all immunopositive for Olig2 in both mouse lines at either age (Extended data Fig. 6-1*A-D*; Fig. 6*B*, *C*), confirming their oligodendrocyte lineage specificity. To analyze the differentiation stage specificity, we immunostained AAV-*TRE*-YFP infected brain slices with an antibody for NG2 or PDGFRα that labels OPCs, or the antibody CC1 that labels post-mitotic oligodendrocytes. The results showed that very few (4-7%) YFP^+^ cells were OPCs, while 92-97% of them were post-mitotic oligodendrocytes in *Plp*-tTA mice 1 or 2 days after virus injection at either age (Fig. 6*B*, *C*). In the case of *Sox10*^+/rtTA^ mice at P14, little difference was observed as only 5-6% of YFP^+^ cells were OPCs 1 or 2 days after virus injection (Fig. 6*B*). However, for *Sox10*^+/rtTA^ mice at P35, approximately 26% of YFP^+^ cells were OPCs 1 day after virus injection (Fig. 6*C*). This observation excluded the possibility that the low OPC percentages in the reporter-containing cells in *Sox10*^+/rtTA^ mice at P14, or in *Plp*-tTA mice at both ages, were due to selective infection by AAV. More intriguingly, the OPC/YFP^+^ cell ratio decreased to 8% after 1 more day in *Sox10*^+/rtTA^ mice at P35 (Fig. 6*C*). These results may suggest that most of the pulse-labeled OPCs in *Sox10*^+/rtTA^ mice at P35 were undergoing terminal differentiation (tOPCs), and *Sox10*^+/rtTA^ increasingly targeted tOPCs from P14 to P35.

**Figure 6.**
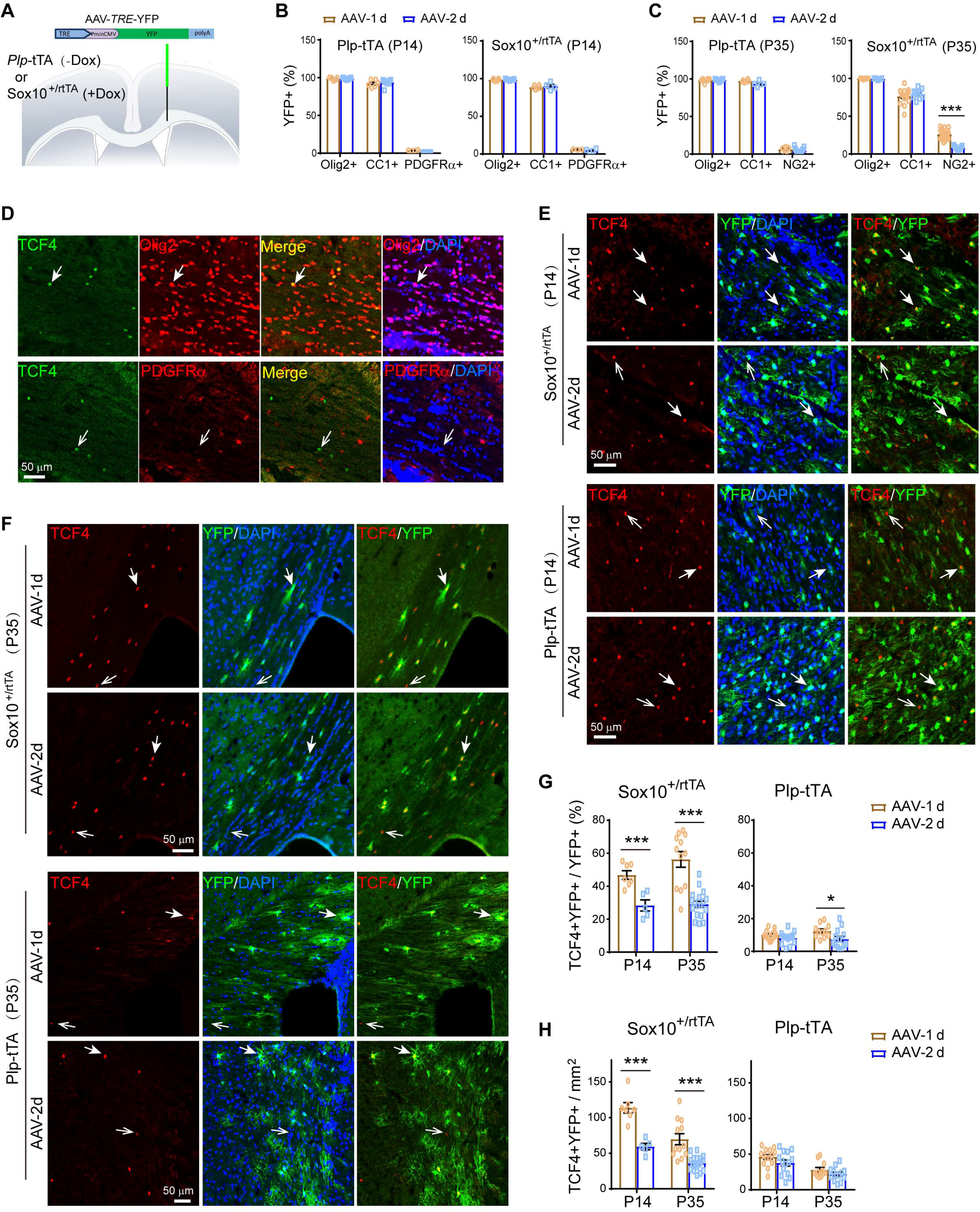
*Plp*-tTA targeted MOs whereas *Sox10*^+/rtTA^ targeted OPC-NFOs in mouse brains during postnatal development. **A,** Schematic illustration of stereotaxic injection sites of AAV-*TRE*-YFP. **B, C,** Percentage of Olig2^+^YFP^+^, CC1^+^YFP^+^, or PDGFRα^+^YFP^+^ (NG2^+^YFP^+^) cells in YFP^+^ cells for *Plp*-tTA or *Sox10*^+/rtTA^ mice 1 (AAV-1d) or 2 days (AAV-2d) after AAV-*TRE*-YFP injection at P14 (**B**) or P35 (**C**). Data were from repeated immunostaining results of 3-7 mice for each group, presented as mean ± s.e.m., and analyzed by unpaired *t* test. For *Plp*-tTA at P14 from AAV-1d to AAV-2d: Olig2^+^YFP^+^ cells, *t*_(12)_ = 0.3698, *P* = 0.718; CC1^+^YFP^+^ cells, *t*_(13)_ = 0.5666, *P* = 0.581; PDGFRα^+^YFP^+^ cells, *t*_(10)_ = 7.532, *P* < 0.0001. For *Sox10*-rtTA at P14 from AAV-1d to AAV-2d: Olig2^+^YFP^+^ cells, *t*_(13)_ = 0.2055, *P* = 0.840; CC1^+^YFP^+^ cells, *t*_(8)_ = 0.6425, *P* = 0.539; PDGFRα^+^YFP^+^ cells, *t*_(5)_ = 1.021, *P* = 0.354. For *Plp*-tTA at P35 from AAV-1d to AAV-2d: Olig2^+^YFP^+^ cells, *t*_(17)_ = 0.4959, *P* = 0.626; CC1^+^YFP^+^ cells, *t*_(9)_ = 2.32, *P* = 0.046; NG2^+^YFP^+^ cells, *t*_(18)_ = 1.003, *P* = 0.329. For *Sox10*-rtTA at P35 from AAV-1d to AAV-2d: Olig2^+^YFP^+^ cells, *t*_(11)_ = 1.098, *P* = 0.296; CC1^+^YFP^+^ cells, *t*_(23)_ = 0.8614, *P* = 0.398; NG2^+^YFP^+^ cells, *t*_(26)_ = 7.869, *P* < 0.0001. **D,** Immunostaining revealed that TCF4 was specifically expressed in a small fraction of Olig2^+^ cells, but not in PDGFRα^+^ cells, in the corpus callosum of mice at P30. Solid arrow, representative double positive cell; Open arrow, representative cell positive for TCF4 only. **E, F,** Double immunostaining results of TCF4 and YFP for brain slices from indicated mice 1 (AAV-1d) or 2 days (AAV-2d) after virus injection at P14 (**E**) or P35 (**F**). Note that TCF4^+^ nuclei in AAV-infected area were almost all localized in YFP^+^ cells in *Sox10*^+/rtTA^ mice 1 day after virus injection (AAV-1d) at either P14 or P35, and the colocalization reduced after 1 more day (AAV-2d). Solid arrows, representative double positive cells; Open arrows, representative cells positive for TCF4 only. **G, H,** The percentage of TCF4^+^YFP^+^ in YFP^+^ cells (**G**), and the density of TCF4^+^YFP^+^ cells (**H**), were analyzed. Data were from repeated immunostaining results of 3-7 mice for each group, presented as mean ± s.e.m., and analyzed by unpaired *t* test. For the percentage in *Plp*-tTA mice from AAV-1d to AAV-2d: at P14, *t*_(26)_ = 1.574, *P* = 0.128; at P35, *t*_(22)_ = 2.367, *P* = 0.027. For the percentage in *Sox10*^+/rtTA^ mice from AAV-1d to AAV-2d: at P14, *t*_(10)_ = 4.39, *P* = 0.0014; at P35, *t*_(28)_ = 6.041, *P* < 0.0001. For the density in *Plp*-tTA mice from AAV-1d to AAV-2d: at P14, *t*_(26)_ = 1.581, *P* = 0.126; at P35, *t*_(22)_ = 1.429, *P* = 0.167. For the density in *Sox10*^+/rtTA^ mice from AAV-1d to AAV-2d: at P14, *t*_(10)_ = 5.685, *P* = 0.0002; at P35, *t*_(28)_ = 4.813, *P* < 0.0001. See also Extended data Figure 6-1.

In most cases from *Sox10*^+/rtTA^ mice, we noticed the total numbers of NG2^+^YFP^+^ (or PDGFRα^+^YFP^+^) and CC1^+^YFP^+^ cells were fewer than that of Olig2^+^YFP^+^ cells (Fig. 6*C*), which indicated the presence of reporter-containing cells that were neither positive for CC1 nor for NG2 (PDGFRα). These cells belong to the NFO stage, which includes both CC1^-^ NG2^-^ (or CC1^-^PDGFRα^-^) and CC1^+^NG2^-^ (or CC1^+^PDGFRα^-^) oligodendrocyte lineage cells. The previous study for *Sox10*^+/rtTA^-targeting specificity only examined the PDGFRα/NG2 and CC1 immunoreactivities in reporter-containing cells. It is known that most NFOs are also positive for CC1 immunostaining (Ye et al., 2009). The β-catenin effector TCF4 is specifically expressed in the NFO stage (Fancy et al., 2009; Fu et al., 2009; Ye et al., 2009), which was present in a subset of Olig2^+^ cells but absent in PDGFRα^+^ cells in mice at P30 (Fig. 6*D*). In *Sox10*^+/rtTA^ mice at P14, immunostaining revealed that approximately 49% of YFP^+^ cells were NFOs (TCF4^+^) 1 day after virus injection, but it reduced to 28% after 1 more day (Fig. 6*E*, *G*). TCF4^+^ cells found in the corpus callosum at P35 were far fewer than P14, and these cells appeared as clusters. Interestingly, in *Sox10*^+/rtTA^ mice at P35, YFP^+^ cells were mostly found in regions with TCF4^+^ cell clusters (Extended data Fig. 6-1*E*), where approximately 56% of YFP^+^ cells were TCF4^+^ 1 day after virus injection and it reduced to 29% after 1 more day (Fig. 6*F*, *G*). The half reduction of NFO percentage in YFP^+^ cells from day 1 to day 2 was consistent with the previous report that NFOs mature into MOs in 1 or 2 days (Xiao et al., 2016). There was another possibility that the transcriptional activity of *Sox10*-rtTA was low in MOs, and thus took more days to have detectable YFP levels. We analyzed the densities of TCF4^+^YFP^+^ cells and found that they reduced to half from day 1 to day 2 after viral infection in *Sox10*^+/rtTA^ mice at either P14 or P35 (Fig. 6*H*). This result excluded the possibility that the reduction of NFO ratio in YFP^+^ cells was due to the increase of targeted MOs, and confirmed the maturation rate of pulse-labeled NFOs from day 1 to day 2. A similar transition rate is applicable for targeted OPCs and NFOs from day 0 to day 1. Through analyses of the stage alteration rate of reporter-containing cells from day 1 to day 2 after viral infection, we can conclude that the *Sox10*^+/rtTA^-targeting cells at day 0 should mostly be OPCs and NFOs.

YFP only have 4 amino-acid residue difference near the chromophore from GFP (Wachter et al., 1998). Therefore, the YFP-containing MOs (CC1^+^TCF4^-^) in *Sox10*^+/rtTA^ mice can be ascribed to the high stability of YFP. We have measured and reported that the half-life of ErbB2 is ∼8 h in HEK293 cells and ∼9 h in breast cancer cells. Moreover, ligand binding stimulates the turnover of ErbB receptors (Liu et al., 2007; Tao et al., 2009; Tao et al., 2014). To clearly examine the oligodendrocyte stages with ErbB2^V664E^ expression, we purified OPCs and induced transgene expression *in vitro*. ErbB2^V664E^ was visualized by immunocytofluorescence in dissociated oligodendrocyte lineage cells from *Sox10*-ErbB2^V664E^ mice, but not those from control mice, with Dox treatment (Fig. 7*A*). Significant ErbB2 immunoreactivities were detected in OPCs (NG2^+^) and NFOs (TCF4^+^) (Fig. 7*B*, open white arrows). Essentially, there were only a few differentiated oligodendrocytes with low MBP levels detected to have ErbB2 signals. Nevertheless, ErbB2 signals in these MBP^+^ (low) cells, which just started maturation, became extremely weak (Fig. 7*B*, solid white arrows). Once oligodendrocytes matured to have high MBP levels, ErbB2 immunoreactivities disappeared (Fig. 7*B*, solid pink arrows), indicating that there were no new ErbB2^V664E^ synthesis and the existed ErbB2^V664E^ degraded during oligodendrocyte maturation. Despite a different oligodendrocyte stage proportion in the *in vitro* system from that *in vivo*, we detected 67.1 ± 8.9 % ErbB2^+^ cells NG2^+^ (OPCs) from the *Sox1*0-ErbB2^V664E^ cultures during early differentiation. Four days *in vitro* after differentiation induction when many MBP^+^ cells were newly developed, 58.7 ± 4.0 % ErbB2^+^ cells were TCF4^+^ (NFOs), and 18.1 ± 2.4 % were MBP^+^ (low) (Fig. 7*C*). There were about 2.92 ± 1.8 % ErbB2^+^ were MBP^+^ (high) where minimal ErbB2 intensities were detected (Fig. 7*C*, *D*). ErbB2 signals were highest in oligodendrocytes at OPC stage, and close to zero at MO stage (Fig. 7*D*), consolidating the concept that ErbB2^V664E^ expression induced by *Sox10*^+/rtTA^ is mainly confined at OPC-NFO stages.

**Figure 7.**
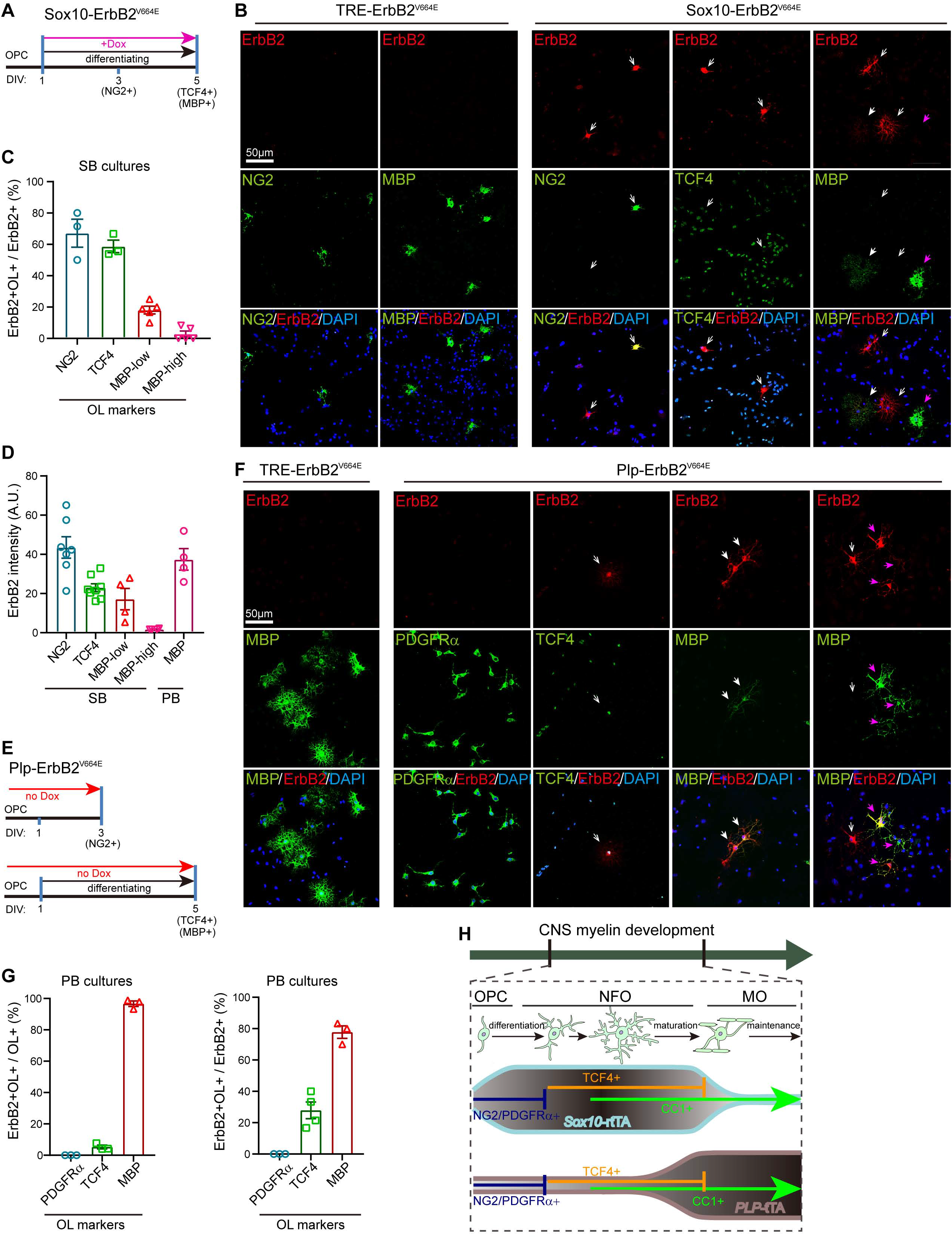
ErbB2^V664E^ mainly expressed in OPC-NFO stages from *Sox10*-ErbB2^V664E^ mice whereas in MOs from *Plp*-ErbB2^V664E^ mice. **A, E,** *In vitro* differentiation and Dox induction protocols for OPCs from *Sox10*-ErbB2^V664E^ (**A**) or *Plp*-ErbB2^V664E^ mice (**E**). DIV, day in vitro. **B, F,** Representative immunostaining images of ErbB2 with protein markers for different oligodendrocyte stages for cultured OPCs from indicated mice with illustrated differentiation and Dox treatment protocols in **A** or **E**. Open white arrows, ErbB2^+^ cells without MBP signals; Solid white arrows, oligodendrocytes with low MBP levels; Solid pink arrows, oligodendrocytes with high MBP levels. **C,** The percentages of ErbB2^+^ cells at different oligodendrocyte stages in total ErbB2^+^ cells detected in each batch staining for cultures from *Sox10*-ErbB2^V664E^ (SB) mice. Note the proportion of oligodendrocytes at each differentiation stage *in vitro* is not the same as that *in vivo*, and ‘Tet-on’ was not switched on for all oligodendrocytes in cultures with Dox supplementation 24 h after cell plating. **D,** The immunostaining intensities of ErbB2 in *Sox10*-ErbB2^V664E^ (SB) oligodendrocytes at each differentiation stage, as well as that in MBP^+^ *Plp*-ErbB2^V664E^ (PB) oligodendrocytes. A.U., absolute unit. **G,** The percentages of ErbB2^+^ cells at different oligodendrocyte stages in total ErbB2^+^ cells detected in each batch staining for cultures from *Plp*-ErbB2^V664E^ (PB) mice, as well as the percentages of ErbB2^+^ oligodendrocytes in total oligodendrocytes at each differentiation stage. ‘Tet-off’ was switched on for all oligodendrocytes in cultures from PB mice, although the proportion of oligodendrocytes at each differentiation stage *in vitro* is not the same as that *in vivo*. **H,** Schematic summary of oligodendrocyte stage-targeting preferences of *Plp*-tTA and *Sox10*^+/rtTA^.

AAV-*TRE*-YFP in *Plp*-tTA mice also labeled some TCF4^+^ cells, which comprised only 7-12% of YFP^+^ cells 1 or 2 days after virus injection at either age (Fig. 6*E*-*H*). It was noticeable that *Plp*-tTA did not specifically target TCF4^+^ cell clusters in the corpus callosum at P35 (Extended data Fig. 6-1*F*). These results implied that *Plp*-tTA did not specifically target the OPC or NFO stage but randomly expressed in the OPC-NFO period at a low ratio. Conversely, 90% of the YFP^+^ cells were TCF4^-^ and 92-97% were CC1^+^ in *Plp*-tTA mice at either P14 or P35, suggesting that *Plp*-tTA steadily targeted MOs. Further, the *in vitro* analyses validated this idea by showing that 96.7 ± 1.7 % MBP^+^ cells (Fig. 7*E-G*), including both cells with low (Fig. 7*F*, solid white arrows) and high MBP levels (Fig. 7*F*, solid pink arrows) from the ‘Tet-off’ *Plp*-ErbB2^V664E^ mice, were ErbB2^+^. There were a few oligodendrocytes not MBP^+^, corresponding to 5.29 ± 1.2 % NFOs (TCF4^+^) with the transgene expression (Fig. 7*F*, open white arrows). None of PDGFRα^+^ cells (OPCs) was immunopositive for ErbB2 (Fig. 7*G*). Note the ErbB2 immunostaining intensities were comparable in NG2^+^ cells from *Sox10*-ErbB2^V664E^ mice and MBP^+^ cells from *Plp*-ErbB2^V664E^ mice (Fig. 7*D*), indicating comparable transactivation efficiencies of *Sox10*-rtTA and *Plp*-rtTA at single cell levels. The *in vivo* and *in vitro* analyses corroborated that *Sox10*^+/rtTA^ and *Plp*-tTA target different oligodendrocyte differentiation stages (Fig. 7*H*), which explains why ErbB overactivation induced distinct cellular and histological phenotypes in *Plp*-ErbB2^V664E^ and *Sox10*-ErbB2^V664E^ mice.

### ErbB overactivation caused MO necroptosis whereas OPC apoptosis

With the advantages of the non-overlapping phenotypes in *Plp*-ErbB2^V664E^ and *Sox10*-ErbB2^V664E^ mice, we were able to investigate the pathogenetic mechanisms that determine the different myelin responses in these two mouse models. MO number reduction occurred earlier than the time when demyelination was obviously observed in the corpus callosum of *Plp*-ErbB2^V664E^ mice, suggesting that primary oligodendropathy may be the cause of demyelination. We noticed there was an increase in the numbers of nuclei associated with CC1^+^ cell debris, starting from 6 dpd, in the corpus callosum of *Plp*-ErbB2^V664E^ mice (Fig. 3A, white arrows). We examined the corpus callosum of *Plp*-ErbB2^V664E^ mice by TdT-mediated dUTP nick end labeling (TUNEL) assay, and observed as little apoptotic signaling as that in controls (Fig. 8*A*-*C*). Immunostaining for cleaved caspase-3, the key apoptotic signaling protein, also exhibited few signals there (Fig. 8*B,C*). These results revealed that the degenerating CC1^+^ cells were necrotic rather than apoptotic. Consistently, the oligodendrocyte nuclei associated with the destroyed myelin sheaths in *Plp*-ErbB2^V664E^ mice were regular nuclei without apoptotic chromatin condensation (Fig. 1*H*, red asterisk). In support of this theory, MLKL, the protein mediating oligodendrocyte necroptosis in multiple sclerosis (Ofengeim et al., 2015) as well as the peripheral myelin breakdown after nerve injury (Ying et al., 2018), demonstrated an increased expression in the callosal CC1^+^ cells in *Plp*-ErbB2^V664E^ mice from 6 dpd (Fig. 8*D*-*F*). Necroptosis is a programmed form of necrosis, and has been revealed to be a prominent pathological hallmark of multiple sclerosis (Ofengeim et al., 2015). RIP3 is the kinase at the upstream of MLKL in this programmed death signaling pathway (Sun et al., 2012; Ofengeim et al., 2015). Notably, the expression of RIP3 was also elevated in the callosal CC1^+^ cells in *Plp*-ErbB2^V664E^ mice from 6 dpd as revealed by both immunostaining and western blotting (Fig. 8*D*-*F*). Significant MLKL signals also appeared in the corpus callosum of adult *Plp*-ErbB2^V664E^ mice at P70 with 10 dpd, indicating similar susceptibility of adult MOs to necroptosis induced by ErbB overactivation (Extended data Fig.3-1*C,E*). Cell necroptosis is a trigger to inflammation (Pasparakis and Vandenabeele, 2015). Based on the timeline, MO necroptosis was the primary defect induced in *Plp*-ErbB2^V664E^ mice, followed by inflammation, OPC regeneration, myelin breakdown, axonal degeneration, and other pathological events as reported in multiple sclerosis (Bradl and Lassmann, 2010; Ofengeim et al., 2015).

**Figure 8.**
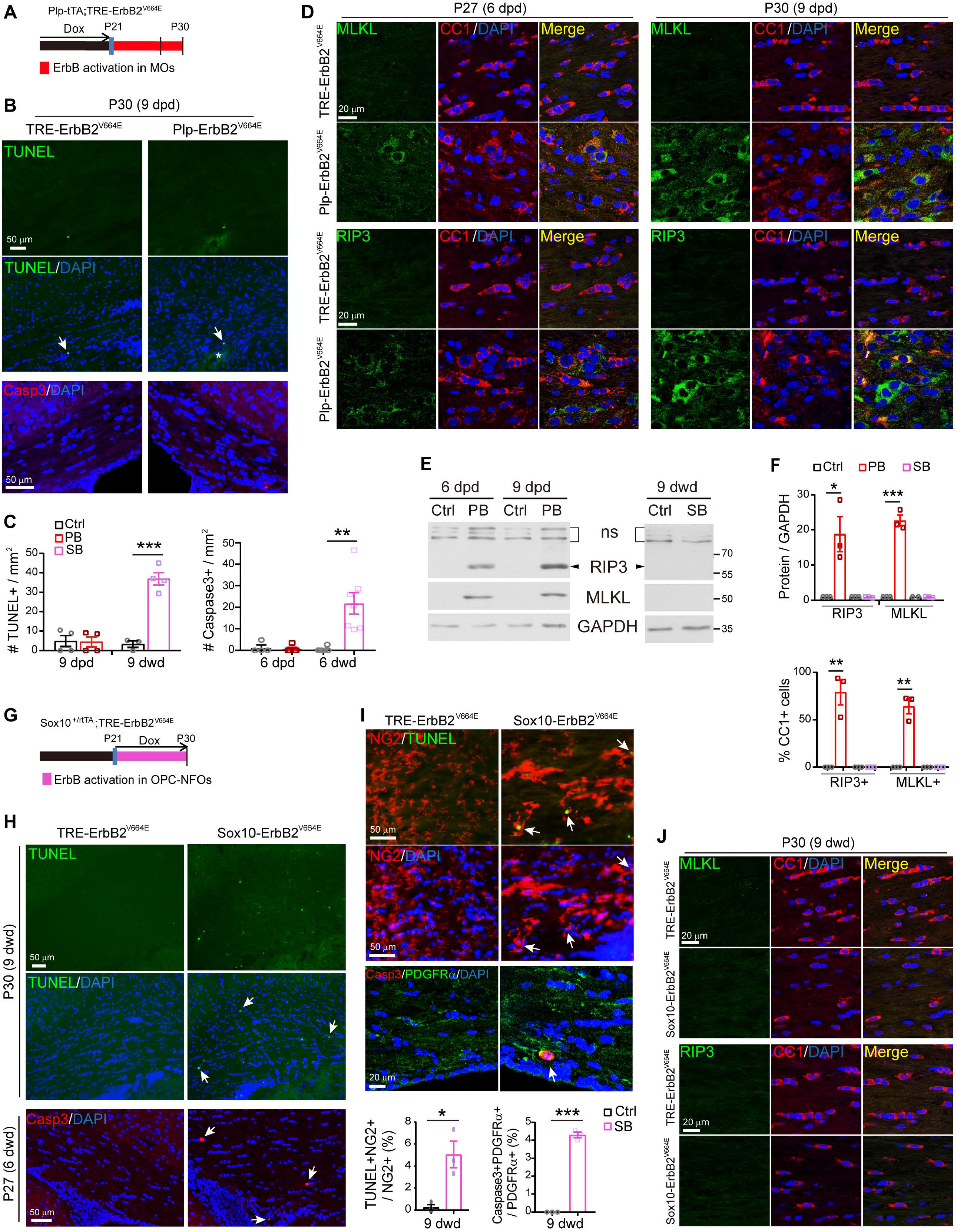
ErbB overactivation induced MO necroptosis in *Plp*-ErbB2^V664E^ mice but OPC apoptosis in *Sox10*-ErbB2^V664E^ mice. **A, G,** Dox treatment setting for indicated mice and littermate controls. **B, H,** Apoptotic cells (arrows) in the corpus callosum of *Plp*-ErbB2^V664E^ and control mice at 9 dpd (**B**), or *Sox10*-ErbB2^V664E^ and control mice at 6 or 9 dwd (**H**), were examined by TUNEL assays and immunostaining for cleaved caspase-3. Note the increased numbers of nuclei (DAPI^+^) and non-specifically stained hemorrhagic spots (asterisk) in the brain slice of *Plp*-ErbB2^V664E^ mice (**B**). **C,** Quantitative data of apoptotic cell densities in indicated mice. PB, *Plp*-ErbB2^V664E^. SB, *Sox10*-ErbB2^V664E^. Ctrl, control mice. Data were presented as mean ± s.e.m., and analyzed by unpaired *t* test. For TUNEL assay: PB, *t*_(6)_ = 0.1128, *P* = 0.914; SB, *t*_(5)_ = 8.344, *P* = 0.0004. For Caspase-3: PB, *t*_(7)_ = 0.3959, *P* = 0.704; SB, *t*_(11)_ = 3.893, *P* = 0.0025. **D, J,** Co-immunostaining results of MLKL or RIP3 with CC1 in the corpus callosum of indicated mice with indicated Dox treatments. **E,** Western blotting results of MLKL and RIP3 in the white matter of *Plp*-ErbB2^V664E^ (PB) mice, or in that of *Sox10*-ErbB2^V664E^ (SB) mice, and in those of littermate control mice (Ctrl). ns, non-specific bands. **F,** Quantitative data of immunostaining and western blotting results of MLKL or RIP3 in indicated mice at 9 days with Dox treatments. Data were presented as mean ± s.e.m., and analyzed by unpaired *t* test. In western blotting results, for RIP3 protein in PB, *t*_(4)_ = 3.579, *P* = 0.023; for MLKL protein in PB, *t*_(4)_ = 13.69, *P* = 0.00017. In immunostaining results, for percentage of RIP3^+^ in CC1^+^ cells in PB, *t*_(4)_ = 6.002, *P* = 0.0039; for percentage of MLKL^+^ in CC1^+^ cells in PB, *t*_(4)_ = 8.202, *P* = 0.0012. **I,** Apoptotic cells (TUNEL^+^ or cleaved caspase-3^+^) were OPCs (NG2^+^ or PDGFRα^+^) in *Sox10*-ErbB2^V664E^ mice at P30 with 9 dwd. Arrows, representative double positive cells. Note OPCs in *Sox10*-ErbB2^V664E^ mice were hypertrophic. The percentage of apoptotic OPCs in total OPCs were quantified and data were presented as mean ± s.e.m. and analyzed by unpaired *t* test. For TUNEL^+^NG2^+^ cells, *t*_(4)_ = 3.95, *P* = 0.0168. For Caspase3^+^PDGFRα^+^ cells, *t*_(4)_ = 26.96, *P* <0.0001.

In contrast, for *Sox10*-ErbB2^V664E^ mice, a dramatic increase in cell apoptosis in the corpus callosum was observed (Fig. 8*C*, *G*, *H*). These apoptotic nuclei, labeled either by TUNEL assay or immunostaining of cleaved caspase-3, were localized in the NG2^+^ and PDGFRα^+^ cells (Fig. 8*I*), indicating apoptosis of OPCs. On the other hand, no increase of MLKL or RIP3 was detected (Fig. 8*E*, *F*, *J*), indicating there was no necroptosis. Developmental apoptosis does not arouse local inflammation, and hence no pathological responses were observed in astrocytes and microglia in the white matter of *Sox10*-ErbB2^V664E^ mice (Fig. 4*J-L*). It is also interesting to note that both the NG2^+^ cells with and without TUNEL^+^ nuclei were hypertrophic in *Sox10*-ErbB2^V664E^ mice (Fig. 8*I*). This phenomenon was not revealed for NG2^+^ cells in *Plp*-ErbB2^V664E^ mice (Fig. 3*B*), further supporting that the OPC phenotypes in *Sox10*-ErbB2^V664E^ mice were attributed to the transgene expression in OPCs.

### Early intervention ceased oligodendrocyte necroptosis/apoptosis and rescued pathological phenotypes in both mice

Next, we explored whether the pathological deficits induced by ErbB overactivation can resolve after correcting the ErbB activities in oligodendrocytes. Toward this end, we withdrew Dox for *Sox10*-ErbB2^V664E^ mice after 9 dwd or re-fed *Plp*-ErbB2^V664E^ mice with Dox after 9 dpd. However, these mice were not rescued. Their disease conditions progressed deleteriously as same as mice with continuous ‘Tet-on’ or ‘Tet-off’. These failures indicated that the pathology in both mice had progressed into an irreversible phase. The delayed effects of Dox feeding or Dox withdrawal on ‘Tet-on’ or ‘Tet-off’ mice may worsen the situations because that the real time for fully suppressing ErbB overactivation was after 9 dwd or 9 dpd for *Sox10*-ErbB2^V664E^ and *Plp*-ErbB2^V664E^ mice. In contrast, *Sox10*-ErbB2^V664E^ mice with 6 dwd, and *Plp*-ErbB2^V664E^ mice with 6 dpd, could be rescued by the Dox-treatment switch. Although both mice at these conditions had exhibited oligodendrocyte necroptosis or apoptosis (Fig. 8*D*, *H*), decreases of myelinating oligodendrocytes (Figs. 3*A*, 5*A*), as well as dysmyelination (Fig. 9*B*, *C*, *I*, *J*), in the corpus callosum on the day before Dox-treatment switching, they became mostly normal after 10-day removal/addition of Dox. The MBP intensities and distributions returned to normal (Fig. 9*A-C*, *H-J*). There were no necroptotic or apoptotic signals in the corpus callosum (Fig. 9*D,K*), and numbers of oligodendrocytes (CC1^+^, Olig2^+^) became as same as those in control mice (Fig. 9*E-G*, *L-N*), despite that proliferative OPCs in *Plp*-ErbB2^V664E^ mice had not completely returned to normal at this time (Fig. 9*E-G*). In a word, MO necroptosis and OPC apoptosis can be ceased by suppressing the aberrant activation of ErbB receptors, and the dead oligodendrocytes, irrespective of inflammatory status, can be cleaned to restore myelin development. These results suggest that the early interference with aberrant ErbB signaling is effective to prevent the white matter pathological progression.

**Figure 9.**
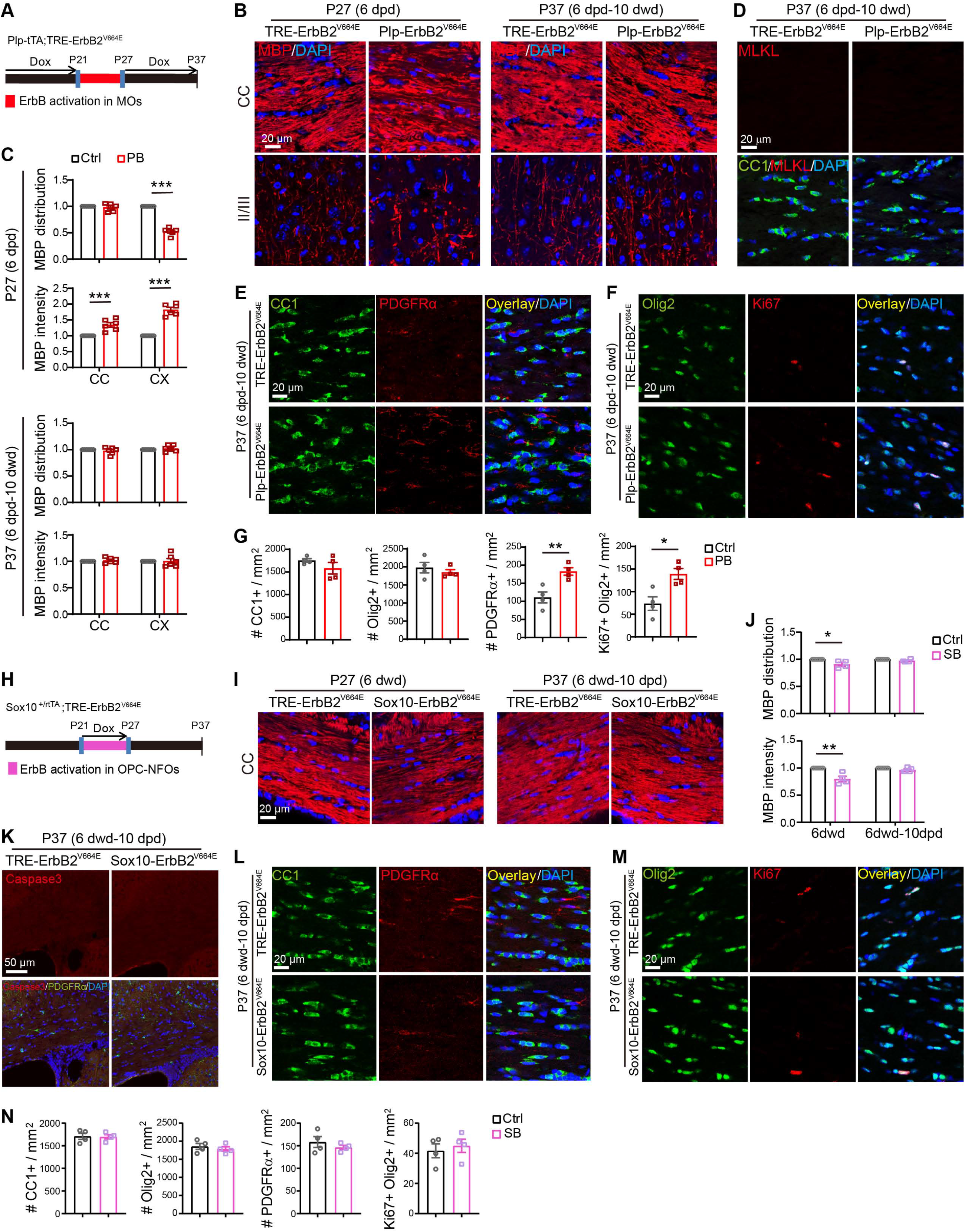
Early intervention rescued pathological phenotypes in both *Plp*-ErbB2^V664E^ and *Sox10*-ErbB2^V664E^ mice. Unless otherwise indicated, quantitative data were presented as mean ± s.e.m., and analyzed by unpaired *t* test. **A, H,** Dox treatment setting for *Plp*-ErbB2^V664E^ (PB) or *Sox10*-ErbB2^V664E^ (SB) mice and littermate controls (Ctrl). **B, I,** MBP immunostaining of indicated mice with indicated Dox treatments. **C, J,** Quantitative data of MBP immunostaining results. Statistical information for PB at P27 (6 dpd) in **C**: MBP distribution of the corpus callosum (CC), *t*_(10)_ = 0.6433, *P* = 0.534; the cortex (CX), *t*_(10)_ = 16.67, *P* <0.0001. MBP intensity of CC, *t*_(10)_ = 4.72, *P* =0.0008; CX, *t*_(10)_ = 11.01, *P* <0.0001. For PB at P37 (6 dpd-10 dwd) in **C**: MBP distribution of CC, *t*_(10)_ = 0.7256, *P* = 0.485; CX, *t*_(10)_ = 0.9144, *P* =0.382. MBP intensity of CC, *t*_(10)_ = 0.8559, *P* = 0.412; CX, *t*_(10)_ = 0.09778, *P* = 0.924. For the corpus callosum of SB in **J**: MBP distribution at P27 (6 dwd), *t*_(7)_ = 2.93, *P* = 0.022; at P37 (6 dwd-10 dpd), *t*_(7)_ = 1.979, *P* =0.088. MBP intensity at P27 (6 dwd), *t*_(7)_ = 4.478, *P* = 0.0029; at P37 (6 dwd-10 dpd), *t*_(8)_ = 1.975, *P* =0.084. **D, K,** No necroptotic (MLKL^+^) or apoptotic signals (cleaved caspase-3^+^) in *Plp*-ErbB2^V664E^ (**D**) and *Sox10*-ErbB2^V664E^ (**K**) mice, respectively, 10 days after Dox treatment switch. **E, F, L, M,** Representative immunostaining results of CC1, PDGFRα, Olig2, Ki67 in the corpus callosum of *Plp*-ErbB2^V664E^ (**E, F**) and *Sox10*-ErbB2^V664E^ (**L, M**) mice 10 days after Dox treatment switch. **G, N,** Quantitative data of oligodendrocyte densities in the corpus callosum. Statistical information for *Plp*-ErbB2^V664E^ (PB) in **G**: for CC1^+^, *t*_(6)_ = 1.288, *P* = 0.245; for Olig2^+^, *t*_(6)_ = 0.7953, *P* = 0.457; for PDGFRα^+^, *t*_(6)_ = 3.907, *P* = 0.0079; for Ki67^+^Olig2^+^, *t*_(6)_ = 3.389, *P* = 0.0147. For *Sox10*-ErbB2^V664E^ (SB) in **N**: for CC1^+^, *t*_(6)_ = 0.1294, *P* = 0.901; for Olig2^+^, *t*_(6)_ = 0.6954, *P* = 0.513; for PDGFRα^+^, *t*_(6)_ = 0.9395, *P* = 0.384; for Ki67^+^Olig2^+^, *t*_(6)_ = 0.5167, *P* = 0.624.

## Discussion

The results from *in vivo* pulse-labeled reporter tracing and *in vitro* transgene-expressing cell examination were consistent with the fact that the death was occurred for OPCs in *Sox10*-ErbB2^V664E^ mice but for MOs in *Plp*-ErbB2^V664E^ mice. These results substantiate that *Sox10*^+/rtTA^ and *Plp*-tTA mice are valuable tools for *in vivo* exploration of molecular and cellular mechanisms that involve oligodendrocyte development and pathogenesis. In the present study, the differentiation-stage targeting preferences of *Sox10*^+/rtTA^ and *Plp*-tTA mice helped manipulate ErbB activities in non-overlapping oligodendrocyte lineage cells, enabling us to distinguish the histological and cellular responses induced by activating ErbB receptors in MOs and OPC-NFOs, respectively. The levels of ErbB2 immunostaining intensities in individual OPCs from *Sox10*-ErbB2^V664E^ mice and individual MOs from *Plp*-ErbB2^V664E^ mice were comparable, suggesting that the transactivation activities of tTA/rtTA at single cell levels induced in *Sox10*^+/rtTA^ and *Plp*-tTA mice are the same (Fig. 7*D*). Moreover, when using real-time RT-PCR to examine the initiation of the transgene in *Sox10*-ErbB2^V664E^ and *Plp*-ErbB2^V664E^ white matter, ErbB2^V664E^ transcript significantly increased in both mice at the first day with Dox addition or removal, suggesting that the *Sox10*-rtTA and *Plp*-tTA have comparable response time to the drug induction. Therefore, the non-overlapping phenotypes observed in *Sox10*-ErbB2^V664E^ and *Plp*-ErbB2^V664E^ mice are attributed to the cellular targeting preferences of *Sox10*^+/rtTA^ and *Plp*-tTA.

By analyzing *Plp*-ErbB2^V664E^ mice, we demonstrated that ErbB overactivation was pathogenetic in MOs through inducing MO necroptosis (Fig. 8*A*-*F*). The inflammatory necroptosis of MOs easily stimulate pathological responses and cytokine releases from glial cells, ependymal cells, endothelial cells, and even possibly infiltrated macrophages in the microenvironment (Burda and Sofroniew, 2014; Pasparakis and Vandenabeele, 2015), leading to a severe pathological complication (Figs. 1-3). Interestingly, studies on genetically modified mice that overexpressed hEGFR in oligodendrocyte lineage cells or overexpressed NRG1 in neurons did not report myelin pathogenesis (Aguirre et al., 2007; Brinkmann et al., 2008). Nevertheless, mice with overactivation of the ErbB downstream signaling in oligodendrocyte lineage cells exhibit myelin and axonal pathology. *Olig2*-Cre;Pten^flox/flox^ mice that overactivate PI3K/Akt signaling in oligodendrocyte lineage cells have loosened myelin lamellae in the spinal cord at 14 weeks and axonal degeneration in the cervical spinal cord fasciculus gracilis at 62 weeks (Harrington et al., 2010). *Plp*-CreER;Mek/Mek mice, which overexpress a constitutively activated MEK, a MAPK kinase, in oligodendrocyte lineage cells with tamoxifen induction, have demyelination in the spinal cord 3 months after the induction of MAPK (Erk) overactivation (Ishii et al., 2016). A dose-dependent detrimental effect is suggested to MEK given the fact that *Plp*-CreER;+/Mek mice exhibit only hypermyelination but not demyelination even 8 months after induction. However, it should be noted that half dose of MEK overactivation also induces astrogliosis and microgliosis, two inflammatory pathological symptoms in the CNS, in the spinal cord 8 months after induction (Ishii et al., 2016), indicating that pathogenetic events might have been sporadically occurred in the white matter of *Plp*-CreER;+/Mek mice. The rapid deterioration in the corpus callosum of *Plp*-ErbB2^V664E^ mice after 9 dpd should be ascribed to significant inflammation that accelerated and aggravated myelin degradation (Fig. 1*L-N*). In contrast, the optic nerve in *Plp*-ErbB2^V664E^ mice had milder inflammatory status and thus had slow myelin degradation and axon degeneration at 14 dpd (Fig. 1*K*, *N*). Mild inflammation may also explain the fact that it takes 3-12 months for *Olig*-cre;Pten^flox/flox^ and *Plp*-creER;Mek/Mek mice to develop myelin breakdown and other pathological changes in white matter (Harrington et al., 2010; Ishii et al., 2016).

Devastating effects of ErbB2^V664E^ in *Plp*-ErbB2^V664E^ mice may be due to its potent promotion of endogenous ErbB activation and multiple downstream signaling including both PI-3K/Akt and MAPK (Erk) pathways (Fig. 1*D*, *E*). There may be a dose-dependent effect for timing or severity of the pathological responses; however, observations in *Plp*-ErbB2^V664E^, *Olig2*-Cre;Pten^flox/flox^, and *Plp*-CreER;Mek/Mek mice corroborate the concept that continuously activating ErbB signaling in oligodendrocyte lineage cells is pathogenetic. Given the fact that *Plp*-CreER and Olig2-Cre mice target the full oligodendrocyte lineage including OPCs (Guo et al., 2009), the cellular mechanism for myelin pathogenesis in the previous reports was not determined. With the advantage of *Sox10*^+/rtTA^ mice that mainly target OPC-NFOs, we demonstrated that overactivation of ErbB receptors that mediate NRG signaling in OPCs induced apoptosis, leading to non-inflammatory hypomyelination (Figs. 4, 5, 8). On the other hand, the study on *Plp*-ErbB2^V664E^ mice suggested that synergistic overactivation of ErbB receptors that mediate NRG (ErbB3/ErbB4) and EGF (EGFR) signaling in white matter leads to more deleterious pathological changes stimulating inflammation (Figs. 1, 3, 8). The differences in *Plp*-ErbB2^V664E^ and *Sox10*-ErbB2^V664E^ mice demonstrated that ErbB overactivation in MOs, but not OPC-NFOs, triggers inflammatory pathogenesis.

The present study revealed that oligodendrocytes and OPCs are cell types susceptible to cell deaths induced by ErbB overactivation, different from astrocytes that become hypertrophic and proliferative with ErbB overactivation (Chen et al., 2017). Notably, it has been reported that overactivation of ErbB receptors can cause cell death, although in many cases ErbB signaling is demonstrated to promote cell proliferation and survival. A number of studies reveal the roles of ErbB4, EGFR, ErbB2 activation in necrosis or apoptosis of macrophages, cancer cells, and a variety of cell types (Hognason et al., 2001; Tikhomirov and Carpenter, 2004; Jackson and Ceresa, 2017; Schumacher et al., 2017). ErbB ligand-induced cell death mostly occurs to cells with high ErbB receptor expression, highlighting that the excessive activation of ErbB receptors is the key to inducing death pathways. Consistently, their downstream Akt and Erk signaling have also been causally linked to cell deaths in some cases (Maddika et al., 2007; Wu et al., 2009; Cagnol and Chambard, 2010). It is intriguing that ErbB receptor overactivation in MOs and OPCs induced distinct deleterious effects. Because that RIP3 and MLKL, the core components of necroptotic machinery, increased specifically in MOs (Fig. 8*D*), but not in astrocytes or microglia which were also pathologically activated in *Plp*-ErbB2^V664E^ white matter (Fig. 1*L*, *M*), the necroptosis signaling should be activated by intracellular signaling pathways downstream of ErbB receptors in MOs. Caspase-8 activation has been reported to be the key event to determine apoptotic fate of cells (Oberst et al., 2011), and defective activation of caspase-8 is critical for RIP1/RIP3/MLKL signaling to induce oligodendrocyte necroptosis in multiple sclerosis (Ofengeim et al., 2015). A cell-type specific RNA-sequencing transcriptome analysis suggests that caspase-8 is minimally expressed in post-mitotic oligodendrocytes but is detectable in OPCs (Zhang et al., 2014), which may determine MO necroptosis but OPC apoptosis under continuous ErbB activation.

The demyelinating disease is now a broad concept that embraces more than multiple sclerosis, neuromyelitis optica, and leukodystrophy. Increasing evidence points to the correlation of the common demyelinating diseases with psychological symptoms (Diaz-Olavarrieta et al., 1999; Chiaravalloti and DeLuca, 2008). On the other hand, white matter lesion is an emerging feature that can be examined periodically by structural brain imaging in patients with psychiatric disorders (Fields, 2008; Mei and Nave, 2014). Remarkably, elevated ErbB activation has been repeatedly implicated in schizophrenia. For example, the mRNA and protein levels of NRG1 and ErbB4 are reported to increase in the prefrontal cortex and hippocampus of schizophrenic patients (Law et al., 2006; Chong et al., 2008; Joshi et al., 2014). These increases could be caused by genetic factors, such as schizophrenia-linked single nucleotide**-**polymorphisms (SNPs) SNP8NRG221132 and SNP8NRG243177, which are identified to increase the mRNA levels of NRG1 (Law et al., 2006). In addition to increased expression, it has been shown that the capacity for ErbB4 to be activated is markedly improved in the prefrontal cortex of schizophrenic patients (Hahn et al., 2006). Besides NRG1 and ErbB4 that have received extensive attention, EGFR is observed to increase in the brain of some schizophrenic patients (Futamura et al., 2002).

The white matter abnormalities observed in our mouse models are reminiscent of diverse myelin-related clinical characteristics in schizophrenic brains, including reduced white matter volume, decreased oligodendrocyte densities, reduced myelin gene products, apoptotic oligodendrocytes, and damaged myelin (Douaud et al., 2007; Uranova et al., 2007; Fields, 2008; Hoistad et al., 2009; Uranova et al., 2011). To our knowledge, we are the first to reveal that ErbB overactivation can primarily induce oligodendropathy and myelin pathogenesis in white matter, providing a possible predisposition of a genetic variability in the ErbB receptors or ligands to the white matter lesion. Notably, SNP8NRG243177 with T-allele, which increases NRG1 Type IV production (Law et al., 2006), is associated with the reduced white matter integrity in schizophrenic patients (McIntosh et al., 2008). The direct evidence provided by the present study that gain of function in ErbB receptors caused primary white matter lesions emphasizes that monitoring white matter in live patients and testing the genetic variability of ErbB signaling pathways are two necessary methods to help develop prognostic and therapeutic strategies for psychiatric disorders.

Intrinsic genetic alterations, apart from the pathogenetic immune responses, may also cause primary oligodendropathy in multiple sclerosis (Factor et al., 2020). Oligodendrocyte necroptosis has been revealed to be a hallmark in multiple sclerosis (Ofengeim et al., 2015), indicating that ErbB receptor activation is possible an upstream pathogenic mechanism for the most common demyelinating disease. It has been reported that inhibition of MAPK (Erk) activity by PD0325901, a MEK inhibitor, promotes OPC differentiation and myelin regeneration in two different animal models of the demyelinating disease (Suo et al., 2019). Moreover, an EGF antibody has been used to neutralize EGF in the animal model of multiple sclerosis and exhibits therapeutic effects by promoting oligodendrogenesis and ameliorating pathological status (Amir-Levy et al., 2014). Therefore, our findings provide a novel molecular and cellular insight into the primary oligodendropathy in demyelinating diseases, and support that early intervention of ErbB overactivation is beneficial in the therapy of the diseases.

## Acknowledgments

We thank Haiping Xiong, Wanwan He, Kaiwei Zhang, and Shasha Zhang in Hangzhou Normal University for the assistance in EM image analyses. This work was supported by grants from the National Natural Science Foundation of China (31371075, 31671070, and 31871030 to YT).

---

**Extended data Figure 1-1.**
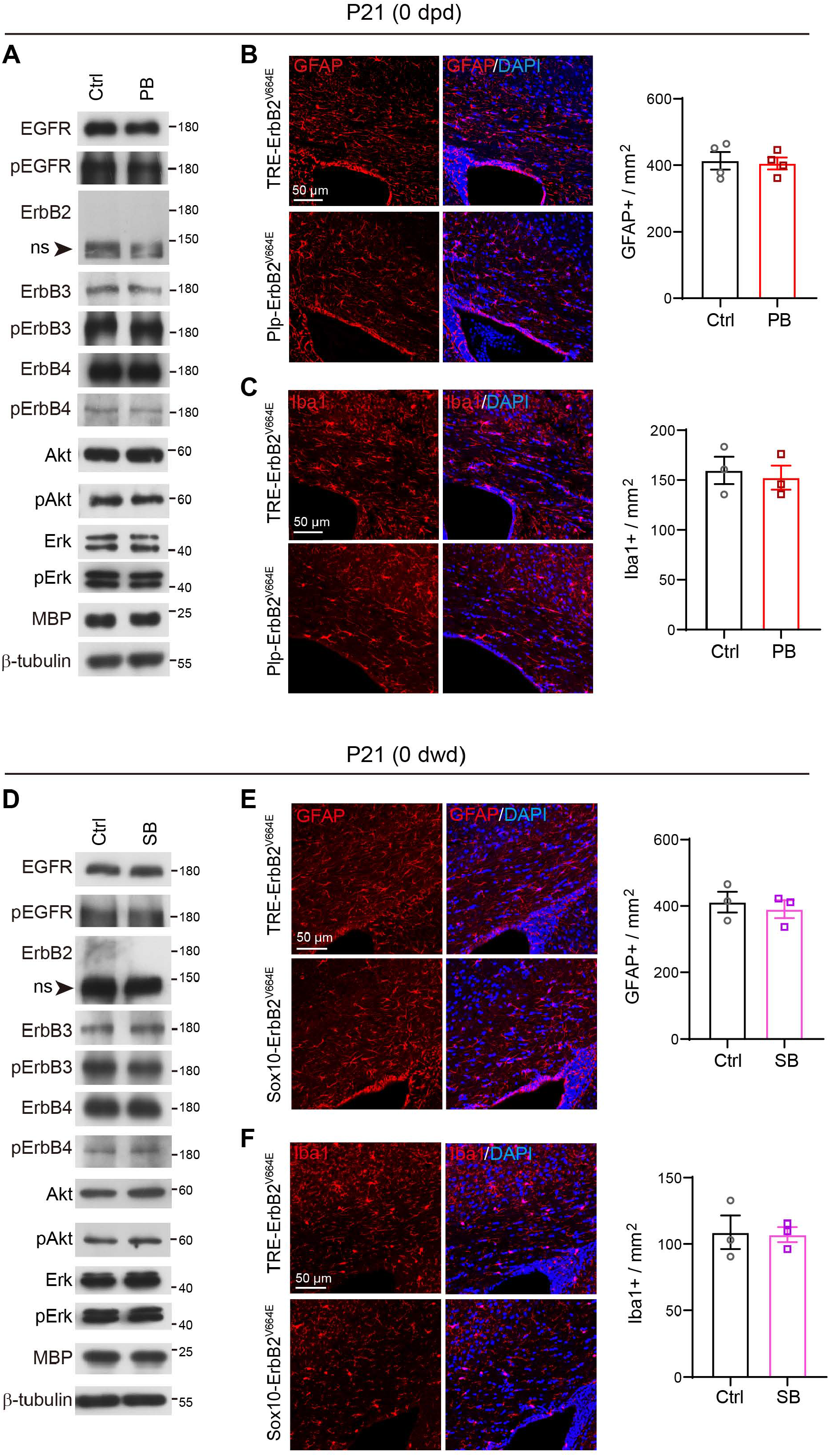
No leak in transgene expression in mice at P21 before drug induction. **A**, **D**, Western blotting of indicated proteins in white matter isolated from *Plp*-ErbB2^V664E^ (PB) mice and littermate control mice (Ctrl) at P21 before Dox withdrawal (**A**), or from *Sox10*-ErbB2^V664E^ (SB) mice and littermate control mice (Ctrl) at P21 before Dox treatment (**D**). ns, non-specific bands. **B**, **C**, **E**, **F**, Astrocytes (GFAP^+^) and microglia (Iba1^+^) examined in the corpus callsoum of indicated mice by immunostaining. Cell densities in the corpus callosum were quantified, and data were presented as mean ± s.e.m.. Statistical analyses by unpaired *t* test revealed no differences.

**Extended data Figure 1-2.**
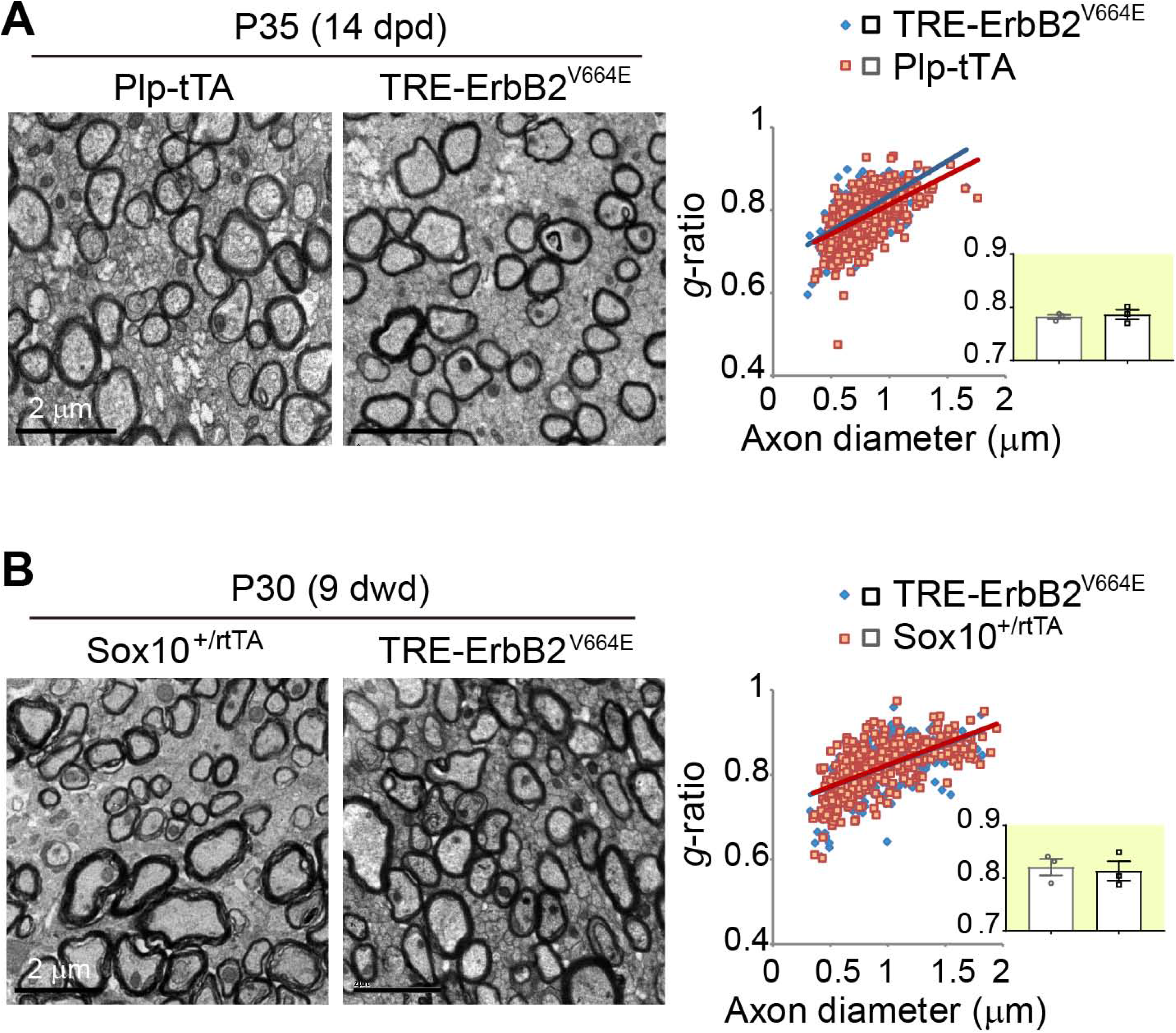
Unaltered myelin in the brains of *Plp*-tTA or *Sox10*^+/rtTA^ mice after Dox treatments. **A, B,** EM images of the corpus callosum of *Plp*-tTA and littermate *TRE*-ErbB2^V664E^ mice at P35 with 14 dpd (**A**), or that of *Sox10*^+/rtTA^ and littermate *TRE*-ErbB2^V664E^ mice at P30 with 9 dwd (**B**). *g*-ratio was calculated for myelinated axons. Averaged *g*-ratio (inset) were presented as mean ± s.e.m., and analyzed by unpaired *t* test. For A, *t*_(4)_ = 0.4472, *P* = 0.678; for B, *t*_(4)_ = 0.3042, *P* = 0.776.

**Extended data Figure 3-1.**
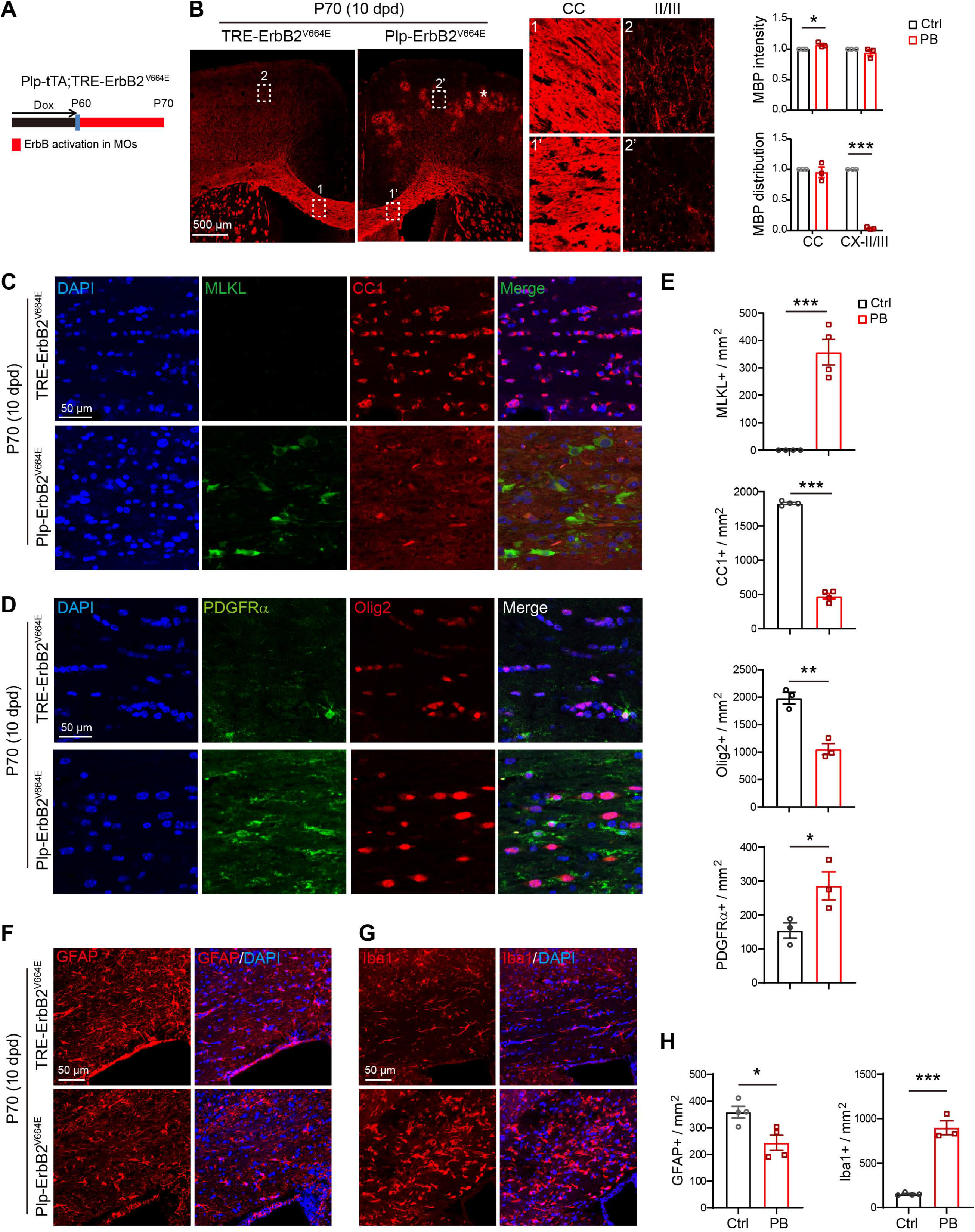
ErbB overactvation induced pathological changes in white matter of adult *Plp*-ErbB2^V664E^ mice. **A,** Dox treatment setting for indicated mice and littermate controls. **B,** MBP immunostaining of *Plp*-ErbB2^V664E^ mice and littermate controls at P70 with 10 dpd. Magnified regions exhibited that myelin structures were comparable in the corpus callosum (CC, 1 and 1’) whereas became fragmented and beaded in layer II/III of the cortex (II/III, 2 and 2’) in *Plp*-ErbB2^V664E^ mice. Note there were more non-specifically stained hemorrhagic spots (white asterisk) in the cortex of these mice than that of *Plp*-ErbB2^V664E^ mice with similar treatment at P30 with 9 dpd. Quantitative data were presented as mean ± s.e.m., and analyzed by unpaired *t* test. For MBP intensity of CC, *t*_(4)_ = 3.033, *P* = 0.0386; CX-II/III, *t*_(4)_ = 1.204, *P* = 0.295. For MBP distribution of CC, *t*_(4)_ = 0.5432, *P* = 0.616; CX-II/III, *t*_(4)_ = 67.58, *P* < 0.0001. **C, D,** Representative immunostaining results of MLKL, CC1, PDGFRα, Olig2 in the corpus callosum of *Plp*-ErbB2^V664E^ and littermate control mice at P70 with 10 dpd. **E,** Quantitative data of oligodendrocyte densities in the corpus callosum were presented as mean ± s.e.m., and analyzed by unpaired *t* test. For MLKL^+^, *t*_(6)_ = 7.716, *P* = 0.0002; for CC1^+^, *t*_(6)_ = 31.34, *P* < 0.0001; for Olig2^+^, *t*_(4)_ = 6.372, *P* = 0.0031; for PDGFRα^+^, *t*_(4)_ = 2.8, *P* = 0.0488. **F**, **G**, Astrocytes (GFAP^+^) and microglia (Iba1^+^) examined in the subcortical white matter of indicated mice by immunostaining. **H**. Astrocyte and microglia densities in the corpus callosum were quantified, and data were presented as mean ± s.e.m., and analyzed by unpaired *t* test. For GFAP^+^, *t*_(6)_ = 3.148, *P* = 0.0199; for Iba1^+^, *t*_(5)_ = 11.36, *P* < 0.0001. Note the GFAP^+^ cell densities were not increased, despite that of Iba1^+^ cells increased dramatically, indicating a different inflammatory profile in adult mice.

**Extended data Figure 4-1.**
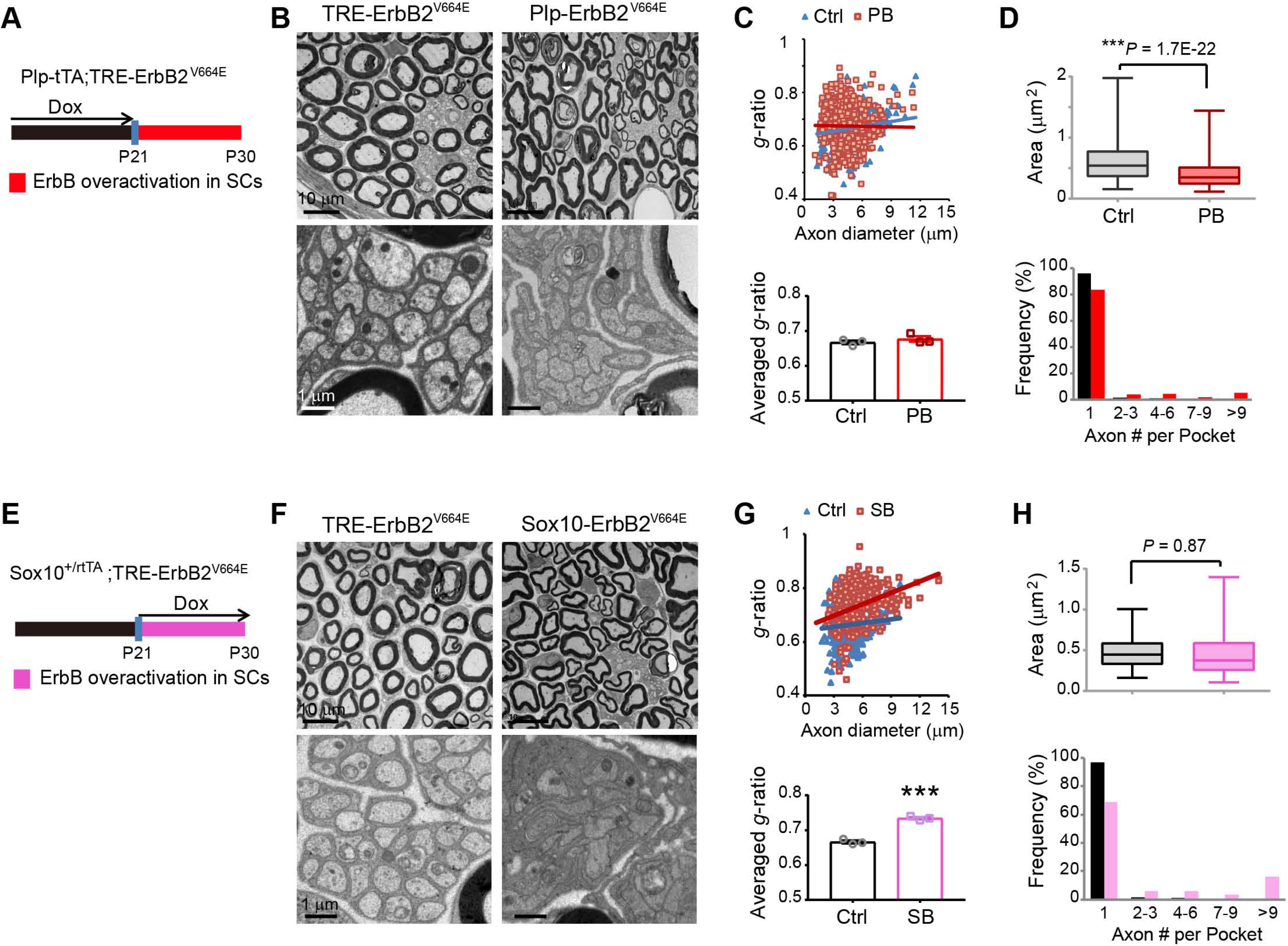
ErbB overactvation induced hypomyelination of peripheral nerves in *Sox10*-ErbB2^V664E^ mice but not in *Plp*-ErbB2^V664E^ mice. **A, E,** Dox treatment setting for indicated mice and littermate controls. SCs, Schwann cells. **B**, **F**, Representative EM images of sciatic nerves for *Plp*-ErbB2^V664E^ with littermate controls (**B**), or *Sox10*-ErbB2^V664E^ with littermate controls (**F**). The top layers of EM images showed myelinated axons in the sciatic nerves, while the bottom layers of EM images showed unmyelinated axons. **C, G,** Quantitative data were shown for *g*-ratio analysis of myelinated axons detected by EM. Averaged *g*-ratio for each mouse were plotted at the bottom, presented as mean ± s.e.m., and analyzed by unpaired *t* test. For **C**, *t*_(4)_ = 1.026, *P* = 0.363; for **G**, *t*_(4)_ = 12.63, *P* = 0.0002. **D, H,** Both *Plp*-ErbB2^V664E^ and *Sox10*-ErbB2^V664E^ mice exhibited slight deficiency in unmyelinated axons of the sciatic nerves. Top: Axonal sizes of unmyelinated axons were measured by their areas in cross sections, and data were plotted as boxes showing quartile and median with whiskers to show 2.5-97.5% of data range. Outlier symbols were omitted. Data were analyzed by unpaired *t* test. Bottom: For the ensheathment analysis of unmyelinated axons, axon numbers in each pocket were counted and quantified by their frequency. For normal mature peripheral nerves, the majority of non-myelinating SC pockets only ensheathe one axon.

**Extended data Figure 5-1.**
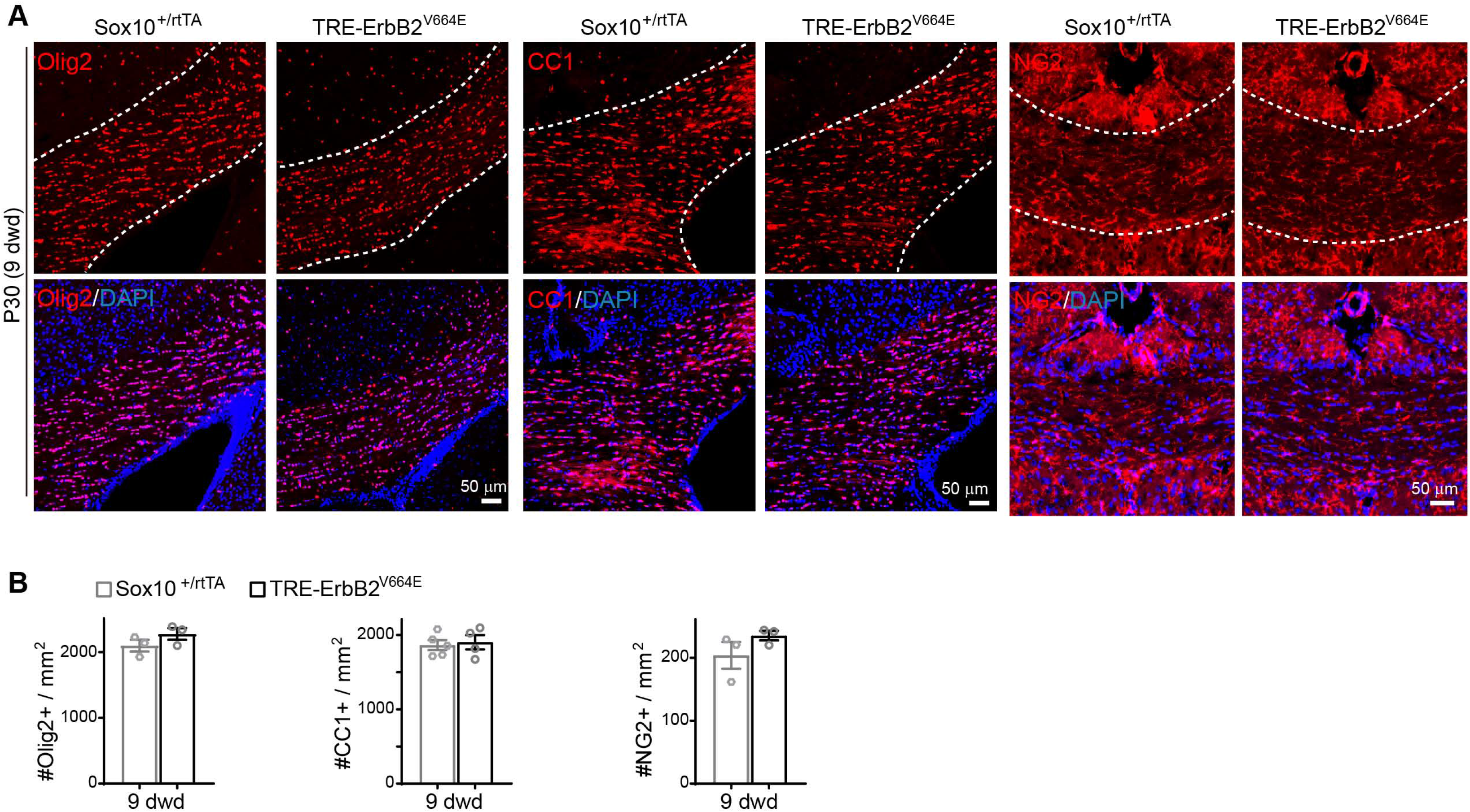
Oligodendrocyte cell densities were similar in the corpus callosum of *Sox10*^+/rtTA^ and littermate *TRE*-ErbB2^V664E^ mice. (**A**) Olig2^+^, CC1^+^, and NG2^+^ cells in the corpus callosum of indicated mice at P30 with 9 dwd were examined by immunostaining. (**B**) Data were from immunostaining of 3 mice for each group, presented as mean ± s.e.m., and analyzed by unpaired *t* test. For Olig2^+^, *t*_(4)_ = 1.418, *P* = 0.229; for CC1^+^, *t*_(7)_ = 0.3431, *P* = 0.742; for NG2^+^, *t*_(4)_ = 1.394, *P* = 0.236.

**Extended data Figure 6-1.**
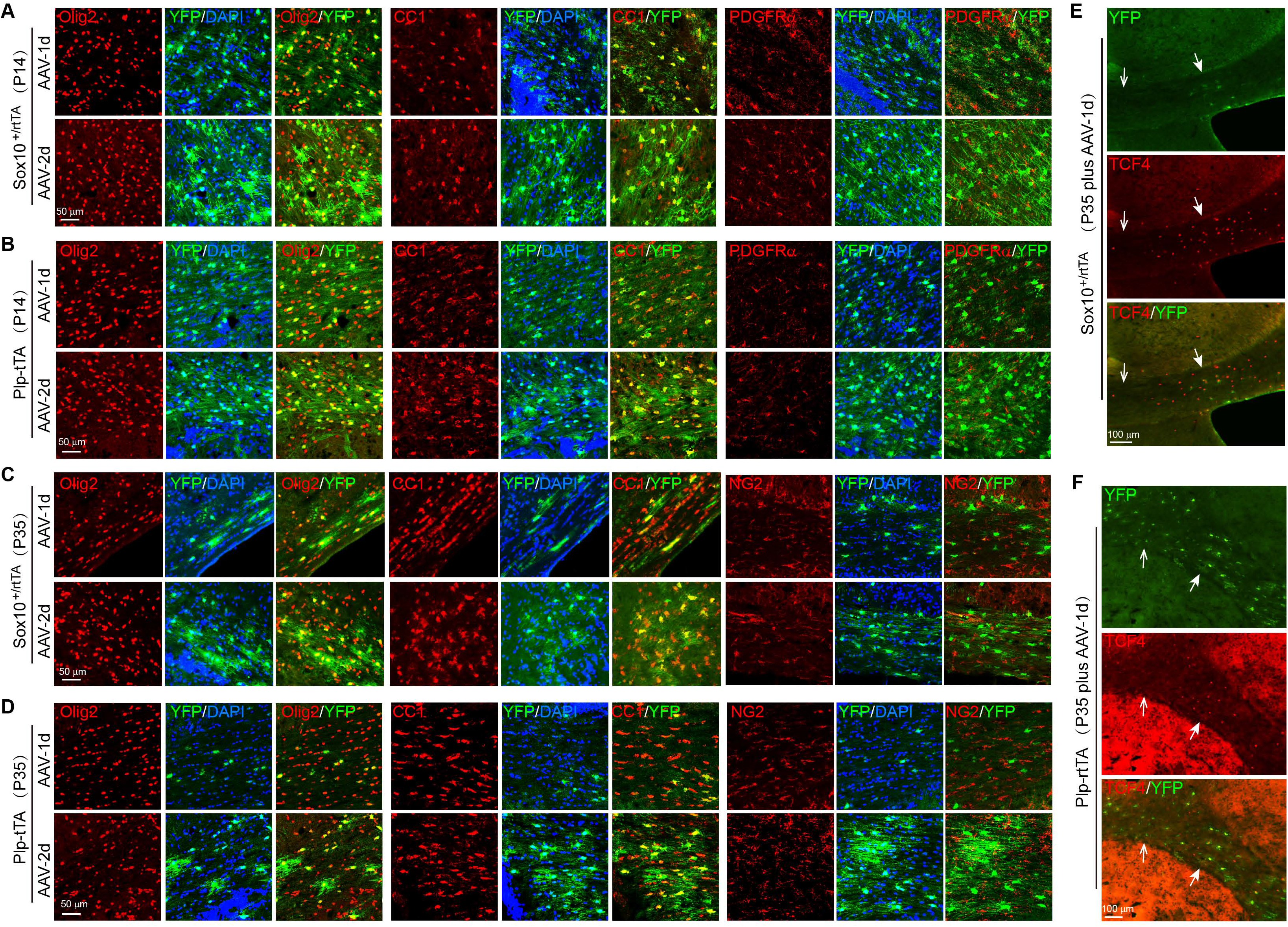
Pulse-labeled reporter-containing oligodendrocytes in *Plp*-tTA and *Sox10*^+/rtTA^ at P14 or P35. **A-D,** AAV-*TRE*-YFP was stereotactically injected into the corpus callosum of *Sox10*^+/rtTA^ or *Plp*-tTA mice at P14 or P35. 1 (AAV-1d) or 2 (AAV-2d) days after virus injection, brain sections were co-immunostained by antibodies to YFP and Olig2, or by CC1 antibody and antibody to YFP, or by antibodies to YFP and NG2 (or PDGFRα). Shown are representative images for indicated mice at P14 or P35. *Sox10*^+/rtTA^ mice were fed with Dox for 3 days before stereotaxic injection of the virus, while *Plp*-tTA mice had no Dox treatment. **E, F,** Distributions of viral pulse-labeled cells, as shown by co-immunostaining of YFP and TCF4, in the corpus callosum of *Sox10*^+/rtTA^ (**E**) or *Plp*-tTA (**F**) mice at P35. Note that reporter-containing (YFP ) cells in *Sox10*^+/rtTA^ mice stringently distributed within TCF4^+^ cell clustered region, whereas those in *Plp*-tTA mice distributed broadly in the corpus callosum. Solid arrows, regions with clustered TCF4^+^ cells; Open arrows, regions with fewer TCF4^+^ cells.

## Notes

**Conflict of interest:** The authors declare no competing interests.

https://doi.org/10.1523/JNEUROSCI.2922-20.2021

### Competing Interest Statement

The authors have declared no competing interest.

